# Cleavage furrow-directed cortical flows bias mechanochemical pathways for PAR polarization in the *C. elegans* germ lineage

**DOI:** 10.1101/2022.12.22.521633

**Authors:** KangBo Ng, Nisha Hirani, Tom Bland, Joana Borrego-Pinto, Nathan W. Goehring

## Abstract

During development, the conserved PAR polarity network is continuously redeployed, requiring that it adapts to changing cellular contexts and environmental cues. How it does so and the degree to which these adaptations reflect changes in its fundamental design principles remain unclear. Here, we investigate the process of PAR polarization within the highly tractable *C. elegans* germline P lineage, which undergoes a series of iterative asymmetric stem cell-like divisions. Compared to the zygote, we observe significant differences in the pattern of polarity emergence, including an inversion of the initial unpolarized state, changes in symmetry breaking cues, and the timings with which anterior and posterior PARs segregate. Beneath these differences, however, polarity establishment remains reliant on the same core pathways identified in the zygote, including conserved roles for cortical actin flows and PAR-dependent self-organization. Intriguingly, we find that cleavage furrow-directed cortical actin flows play a similar symmetry-breaking role for the germline cell P1 as centrosome-induced cortical flows in the zygote. Through their ability to induce asymmetric accumulation of PAR-3 clusters, these furrow-directed flows directly couple the geometry of polarization to cell division, which could be a general strategy for cells to ensure proper organization within dynamically growing systems, such as embryos. In summary, our data suggest that coupling of novel symmetry-breaking cues with highly adaptable core mechanochemical circuits enable robust PAR polarity in response to changing developmental contexts.

## Introduction

During development, the highly conserved PAR polarity network is continuously redeployed in a variety of cell and tissue types. However, how the network adapts and responds to changing cellular contexts and environmental cues remains poorly understood. Indeed, differences in how polarity emerges in different cells often make it difficult to generalize principles.

As a model for understanding how the PAR network behavior adapts during development, we used the highly tractable *C. elegans* germline P lineage. The germline precursors in *C. elegans* are set aside during a program of four sequential PAR-dependent asymmetric stem cell-like divisions. Beginning with P0 and proceeding through P1, P2 and P3, each cell polarizes and undergoes an asymmetric division that yields one P lineage daughter, which retains germline potential (Figure 1A). These divisions ultimately give rise to the germline founder cell, P4, which divides symmetrically to generate the two primordial germ cells, Z2 and Z3 (Rose and Gönczy, 2014; Sulston et al., 1983).

**Figure 1.**
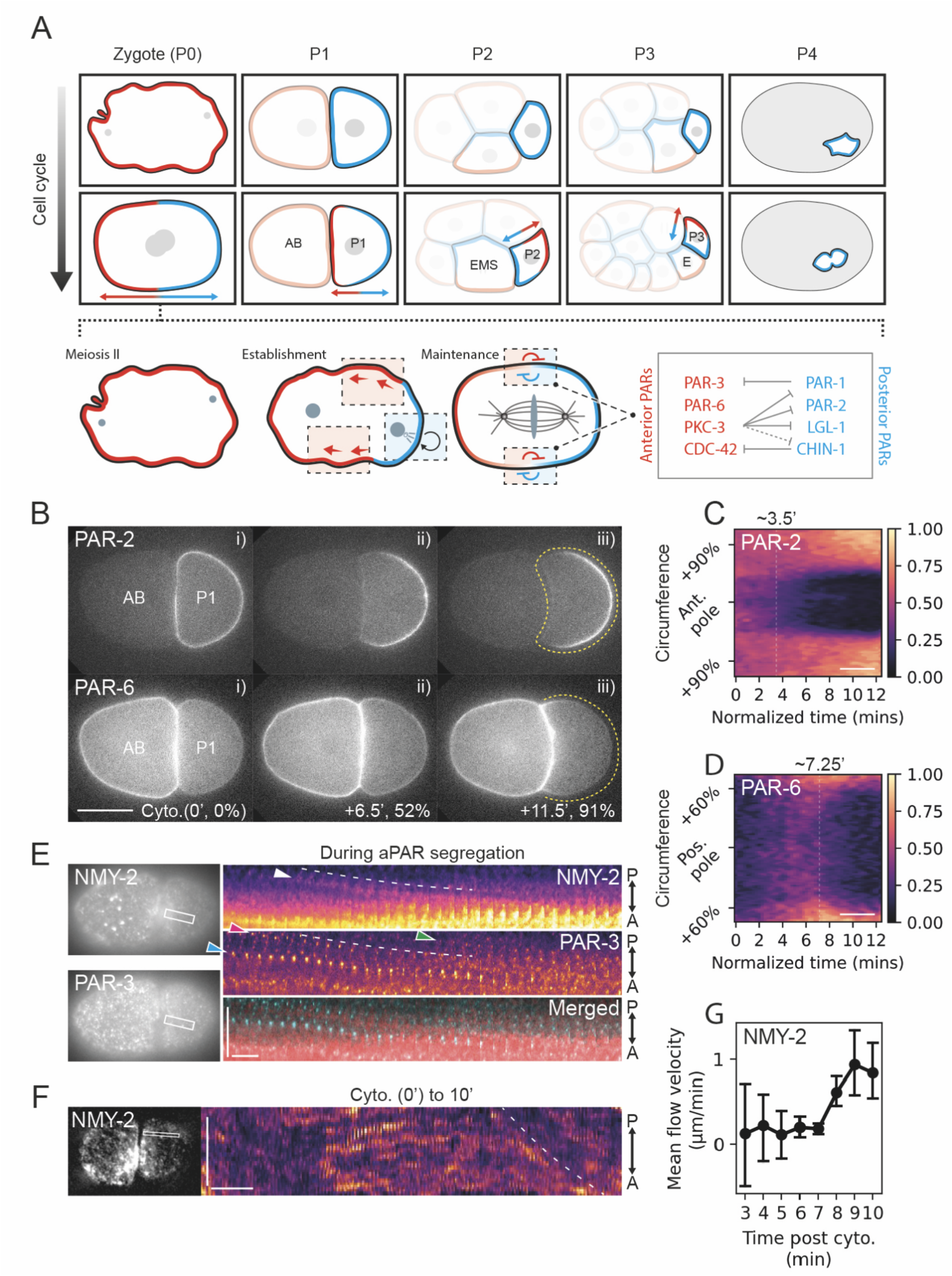
Polarization of pPARs precede aPAR segregation during P1 polarity establishment. (A) Iterative polarization of PAR proteins is associated with asymmetric division of P lineage blastomeres. Top, a schematic of PAR localizations in each blastomere at early and late stages of the cell cycle. Note the reversal of PAR domain orientation in P2 and P3 such that the PAR-2 domain (cyan) is oriented towards the E lineage neighbor (EMS, E). Below - for reference, the general mechanisms of polarization as understood from work in the zygote. Here, prior to symmetry breaking, the membrane is uniformly enriched with aPARs (red) and depleted of pPARs (cyan). The centrosome then induces symmetry breaking by triggering cortical flows (dotted red box), which segregates aPARs to the anterior, and induces self-organization of a PAR-2 domain (dotted blue box) at the posterior through microtubule binding. Once polarity has been established, mutual antagonism (dotted box with gradient) keeps aPAR and pPAR domains segregated. Mutual antagonism is driven by (i) phosphorylation of PAR-3 by PAR-1, which inhibits clustering, (ii) deactivation of CDC-42 by CHIN-1, and (iii) phosphorylation-dependent membrane displacement of PAR-1, PAR-2, and LGL-1 by PKC-3. Anterior exclusion of CHIN-1 is PKC-3 dependent, but it is still unknown whether CHIN-1 is a direct target. (B) A time series of midsection confocal images of embryos expressing both mCherry::PAR-2 and PAR-6::mNG (NWG0268) during polarity establishment in P1. Note that posterior segregation of PAR-2 precedes segregation of PAR-6 into the anterior. Yellow dotted lines indicate the region used for membrane quantification in (C) and (D). PAR-6 at cell contacts (anterior) was not quantified, as fluorescence signals there could be obscured by signals from AB. Scale bar, 20μm. Time is indicated in minutes, and % interval between cytokinesis and NEBD. (C, D) Averaged spatiotemporal profile of PAR-2 (C) and PAR-6 (D) membrane levels from completion of zygote cytokinesis to nuclear envelope breakdown (NEBD) (n=6). Scale bar, 3 mins. Time of individual movies was normalized to the interval between cytokinesis and NEBD (12.6 ± 0.34 mins). Time (mins) indicated above plot indicates mean non-normalized time of PAR segregation. (E) Representative cortical images and kymograph of mNG::PAR-3 and NMY-2::mKate (NWG0150) during aPAR segregation in P1 (n=6) showing co-movement of PAR-3 and NMY-2 towards the anterior (bottom). White box indicates the region of the kymograph. (Top) White arrowhead indicates movement of a myosin spot towards the anterior. (Bottom) Green, magenta and cyan arrowheads indicate three separate PAR-3 puncta moving towards the anterior. Horizontal scale bar: 10s, vertical scale bar: 10μm. (F) Representative cortical images and kymograph of NMY-2::GFP (LP162). Horizontal scale bar: 1 min, vertical scale bar: 10μm. Dashed line indicates onset of flows. (G) Quantification of PIV analysis of NMY-2 flow velocity at indicated times (n=5). Mean and 95% confidence intervals (bootstrapped) are shown.

Most of what we know about PAR polarity in this system comes from study of the zygote (P0). Here, PAR polarity relies on mutual antagonistic cross-talk between two groups of proteins that form opposing membrane-associated domains (Figure 1A): anterior or aPAR proteins (PAR-3, PAR-6, PKC-3, CDC-42) and posterior or pPAR proteins (PAR-1, PAR-2, LGL-1, CHIN-1) (Beatty et al., 2010; Boyd et al., 1996; Etemad-Moghadam et al., 1995; Gotta et al., 2001; Guo and Kemphues, 1995; Hoege et al., 2010; Kumfer et al., 2010; Tabuse et al., 1998; Watts et al., 1996). Each group excludes the other from its respective domain: the polarity kinase PKC-3 excludes pPARs from the anterior, while the kinase PAR-1 and the CDC-42 GAP CHIN-1 suppress aPARs in the posterior (Kumfer et al., 2010; Motegi et al., 2011; Sailer et al., 2015).

Prior to symmetry breaking, aPARs are enriched uniformly throughout the plasma membrane and largely restrict pPARs to the cytoplasm (Cowan and Hyman, 2004; Schonegg and Hyman, 2006). Symmetry is broken through two semi-redundant cell-intrinsic cues (Figure 1A). First, cortical actin flows, which typically manifest alongside pronounced ruffling of the cell cortex, segregate aPARs towards the nascent anterior half of the cell and allow pPARs to associate with the posterior membrane (Goehring et al., 2011b; Munro et al., 2004). Second, formation of the pPAR domain is further promoted by the ability of PAR-2 to self-organize (Arata et al., 2016; Hao et al., 2006; Motegi et al., 2011). This so-called PAR-2 pathway relies on the combined effects of the RING domain and microtubule-binding activity of PAR-2, which provides membrane stabilization and local protection from PKC-3 phosphorylation at the posterior respectively (Hao et al., 2006; Motegi et al., 2011). As a nascent pPAR domain forms, PAR-2 stabilizes PAR-1, which in turn promotes local displacement of aPARs through phosphorylation of PAR-3 (Motegi et al., 2011). While cortical flows appear to be the dominant early cue in wild type zygotes, when flows are absent or compromised, the PAR-2 pathway is sufficient for polarization, allowing pPARs to invade uniform aPAR-enriched membranes to form a domain that eventually gains the capacity to exclude aPARs (Gross et al., 2019; Motegi et al., 2011; Zonies et al., 2010).

Previous work has shown that asymmetric divisions of later P lineage cells retain a dependence on PKC-3 for polarization, consistent with a continued requirement for PAR polarity (Hubatsch et al., 2019). However, the extent to which polarity pathways found in P0 are redeployed in later P lineage cells, and whether individual PAR proteins remain involved is largely unexplored. Several observations suggest that the machinery driving polarization may differ. First, because the P lineage daughter necessarily arises from the pPAR-enriched half of the cell during asymmetric division, P1-P3 are all born with a cortex enriched for pPAR proteins, in striking contrast to P0, in which the cortex is initially aPAR enriched (Figure 1A) (Hubatsch et al., 2019). Second, polarity in P1-P3 is not associated with large scale membrane ruffling associated with cortical actin flows in P0 (Munro et al., 2004). Third, experiments using a temperature sensitive *par-2* allele suggest that the requirement for PAR-2 may be lost in later P lineage divisions (Cheng et al., 1995). Finally, P2 and P3 exhibit a so-called polarity reversal, consistent with a switch from intrinsic to extrinsic polarity cues, which is critical for ensuring the proper orientation of polarity and asymmetric divisions with respect to neighboring cells (Arata et al., 2010; Schierenberg, 1987). In P2 and P3, signaling with their respective endoderm precursor neighbors, EMS and E, promote segregation of PAR-2 towards, rather than away from the cell contact, which occurs in P1 (Figure 1A). Altogether, these data imply substantial modification of polarity pathways in these closely related cells.

Here we sought to characterize the mechanisms of polarization in the P lineage with a focus on P1. Specifically, we asked if the polarity pathways found in the zygote are also deployed in the P lineage or alternatively, if PAR polarization in later P lineage cells represent a shift in paradigm. We found that despite a visually distinct pattern of polarity establishment, the pathways used for polarization appear highly conserved in P lineage blastomeres, including roles for PAR-2 self-organization, mutual antagonism between aPARs and pPARs, and cortical flows. Notably, we find a conserved role for symmetry-breaking cortical flows, which in P1 are directed by the ingressing cleavage furrow rather than the centrosome as in the zygote. By concentrating PAR-3 at the nascent cell contact, these furrow-directed flows effectively couple daughter cell polarity to the prior cell division. Thus, our work both highlights (1) the intrinsic adaptability of core polarization mechanisms to changing cellular contexts and symmetry-breaking cues, and (2) suggests a role of cleavage furrow-directed cortical flows in linking the geometry of PAR polarity to cell division, which could be a broadly applicable strategy for ensuring proper cell organization in dynamically growing systems.

## Results

### pPAR polarization precedes aPAR segregation in P1

To understand how PAR polarity is reestablished in P lineage blastomeres, we first characterized the behavior of aPARs and pPARs through the cell cycle of the more tractable P1 cell. By imaging P1 immediately following division of the zygote, we found that pPARs initially appeared uniformly enriched at the membrane, while aPARs were either confined to the cytoplasm or present very weakly at the plasma membrane (Figure 1B-i, Figure S1). This initial state of P1 is inverse to that of zygote, in which aPARs are initially enriched at the plasma membrane while pPARs are cytoplasmic (Figure 1A, B, Figure S1).

The first obvious signs of polarization in P1 typically occurred within 3-5 minutes following completion of cytokinesis in the zygote, as PAR-2 cleared from region of the cell contact with AB, which corresponds to the anterior of P1, and was concentrated into a well-defined posterior domain (Figure 1B-ii, C, Figure S1A, B). Importantly, we cannot rule out some level of pre-existing aPAR asymmetry in P1, due to high membrane concentration of aPARs in AB and the coincidence of AB and P1 membranes at the cell contact site.

Surprisingly, despite the formation of a PAR-2 domain, aPARs loaded onto the plasma membrane in a relatively uniform manner, such that aPAR and pPARs overlapped in the posterior (Figure 1B-ii, Figure S1C-F). Thus, PAR-2 appears to self-organize into a posterior domain prior to segregation of aPARs. This behavior is similar to what has been observed in the zygote when the predominant symmetry-breaking cue, actin flow, is disrupted (Gross et al., 2019; Motegi et al., 2011; Zonies et al., 2010). Only after approximately 7 mins, did aPARs begin to segregate from the posterior (Figure 1B-iii, D). Finally, by ∼10 mins, P1 exhibited the typical complementary, mutually-exclusive arrangement of aPAR and pPAR domains (Figure 1B-iii, Movie S1).

Thus, the process of PAR polarization in P1 appears to be inverted relative to the zygote: (i) pPARs are initially enriched at the plasma membrane, while aPARs are predominantly cytoplasmic; (ii) an early symmetry-breaking cue induces segregation of PAR-2; (iii) aPARs subsequently load and segregate away from the posterior, following formation of a pPAR domain (compare this sequence to Figure 1A - Zygote).

### Actin flow and pPAR antagonism contributes to segregation of aPARs in P1

In the zygote, cortical actin flows and posterior exclusion by pPARs act in a semi-redundant manner to drive aPAR segregation towards the anterior (Motegi et al., 2011). Munro et al. (2004) reported contemporaneous flows of PAR-6 and NMY-2 to the anterior of P1. Consistent with these findings, during aPAR segregation, PAR-3 and NMY-2 appeared to flow together towards the cell contact (cell anterior) (Figure 1E). Moreover, we observe NMY-2 flow velocities undergoing a sharp increase at the time of onset of aPAR segregation (Figure 1F, G). Note that in contrast to the symmetry-breaking flows in the zygote, these flows occur well after PAR-2 segregation, and therefore we refer to them as “late flows.”

To determine whether these “late flows” are necessary and/or sufficient for aPAR segregation in P1, and whether they act redundantly with pPAR exclusion of aPARs, we inhibited cortical flows and pPAR-dependent exclusion alone or in combination. To eliminate cortical flows, we utilized a fast-acting temperature sensitive allele of *nmy-2* (Liu et al., 2010), and shifted the temperature three minutes after cytokinesis (Figure S2A), which was the earliest time that did not lead to regression of the cleavage furrow (data not shown). To disrupt pPAR-dependent exclusion we utilized an allele of *par-3, par-3(S950A*), which lacks the dominant PAR-1 phosphorylation site (Motegi et al., 2011). Importantly, because of the redundancy of the two pathways in the zygote, *par-3(S950A)* zygotes still polarize (Figure S3). The latter was critical, as it allowed us to use *nmy-2(ts*); *par-3(S950A)* zygotes to generate P1 cells at the permissive temperature and then inactivate cortical flows through acutely shifting 2-cell embryos to the restrictive temperature (Figure S3).

Regardless of whether one or both pathways was inhibited in P1, PAR-2 still segregated into a posterior domain, which is similar to what has been observed in zygotes (Figure 2A, B) (Motegi et al., 2011). By contrast and similar to the zygote, while disruption of either pathway alone had no obvious effect on aPAR segregation, aPARs remained uniformly enriched throughout the plasma membrane when both pathways were inhibited (Figure 2B, C). Taken together, our data suggests that similar to the zygote, both cortical flows and pPAR-dependent aPAR exclusion from the posterior contributes to aPAR segregation in P1.

**Figure 2.**
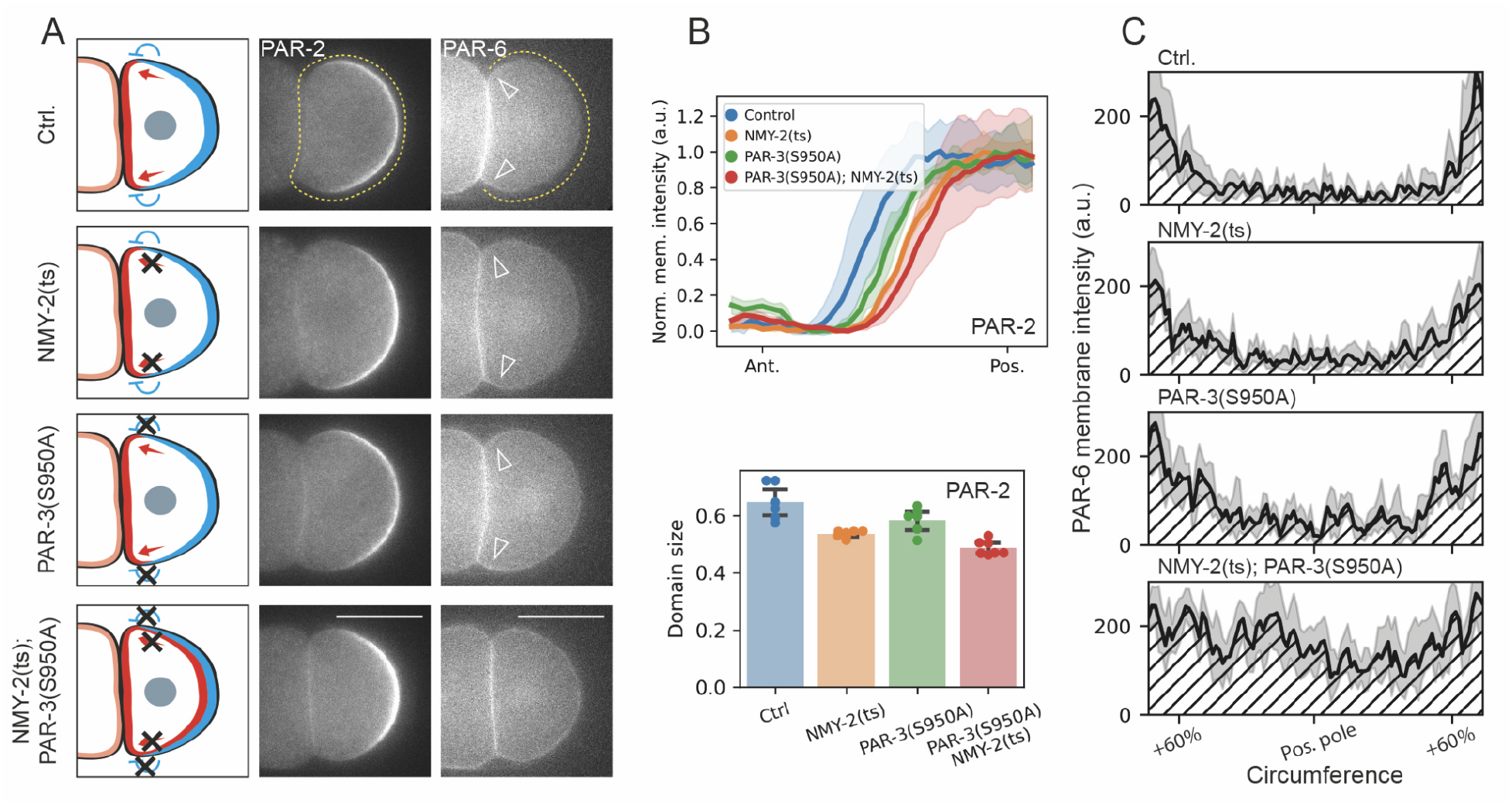
Actin flows and pPAR antagonism act redundantly to polarize aPARs in P1. (A) Schematic of proposed mechanisms of aPAR segregation shown at left (flows, red arrows; aPAR exclusion, blue inhibitory arrow). The schematic indicates the blocked mechanisms and its corresponding phenotypes in P1. At right, representative midsection confocal images of GFP::PAR-2 and PAR-6::mCherry in a wild-type (Ctrl.; NWG0076) (n=6), *nmy-2(ts)* (NWG0283) (n=7), *par-3(S950A*) (NWG0258) (n=6), or a double *nmy-2(ts)*; *par-3(S950A*) background (NWG0319) (n=7) at NEBD. Note that PAR-6 in *nmy-2(ts); par-3(S950A*) embryos fail to segregate from the posterior. Open arrowheads indicate segregation of PAR-6 towards the anterior. Scale bar, 20μm. (B) Quantification of PAR-2 membrane distribution and domain size indicating that the corresponding genotypes cause changes in domain size, but not its ability to polarize. PAR-2 membrane profiles extracted from around the full P1 circumference as indicated in (A, PAR-2). (C) Quantification of PAR-6 membrane distribution extracted from around the non-contact membrane as indicated in (A, PAR-6). Mean and 95% confidence interval (bootstrapped) indicated.

### The PAR-2 pathway promotes early polarization of pPARs in P1

We next sought to determine whether the polarization of pPARs in P1 relies on the same pathways that operate in the zygote. In wild type P1 cells, pPAR polarization takes place prior to segregation of aPARs from the posterior (Figure 1B-ii), and as we have shown can proceed in the absence of aPAR segregation (Figure 2A, B). Previous work has shown that polarization of pPARs in the zygote can also proceed in the absence of aPAR segregation (Motegi et al., 2011). However, this result does not imply that aPAR activity is not required as either depletion or inhibition of PKC-3 completely blocks pPAR polarization in the zygote (Hao et al., 2006; Motegi et al., 2011; Ng et al., 2022; Rodriguez et al., 2017). To determine whether the same was the case in P1, we turned to an analog sensitive version of PKC-3, PKC-3^AS^. Consistent with previous findings, and in support of a strict requirement for PKC-3 activity in P1, acute treatment of *pkc-3*^*AS*^ embryos with the ATP analog 1NA-PP1 at birth of P1 completely blocked pPAR polarization (Figure S4) (Hubatsch et al., 2019; Ng et al., 2022).

In the zygote, the ability of pPARs to polarize in the absence of aPAR segregation is thought to rely on PAR-2 self-organization in response to local microtubule cues provided by the centrosome (Motegi et al., 2011). It has been postulated that the ability for PAR-2 to bind to microtubules provide local resistance to PKC-3 phosphorylation, and enables PAR-2 to load onto the posterior membrane in presence of aPARs. Subsequently, the RING domain of PAR-2 helps stabilize the protein on the posterior membrane even after the centrosome departs from the posterior membrane (Hao et al., 2006; Motegi et al., 2011). Thus, in absence of aPAR segregation, pPAR polarization in the zygote becomes sensitive to mutations in either the PAR-2 RING domain, or the PAR-2 microtubule binding sites (Gross et al., 2019; Hao et al., 2006; Motegi et al., 2011). We therefore sought to determine whether these requirements were also maintained in P1.

To selectively disrupt microtubule binding of PAR-2, we introduced the previously described R183-5A mutation into the endogenous locus, hereafter *par-2(MT-)* (Motegi et al., 2011). Because polarization is largely unaffected in *par-2(MT-)* zygotes when cortical flows are normal, they undergo normal asymmetric division to generate AB and P1 (Figure S5A). Although PAR-2(MT-) supported a smaller PAR-2 domain (Figure 3A, B), which we also observed in the zygote (Figure S5A), counter-intuitively, it polarized to the posterior even earlier than PAR-2(WT) (Figure 3A, C). This result suggests that microtubule binding by PAR-2 at the posterior is not the dominant symmetry breaking cue in P1 (Figure 3A-C, Figure S5E).

**Figure 3.**
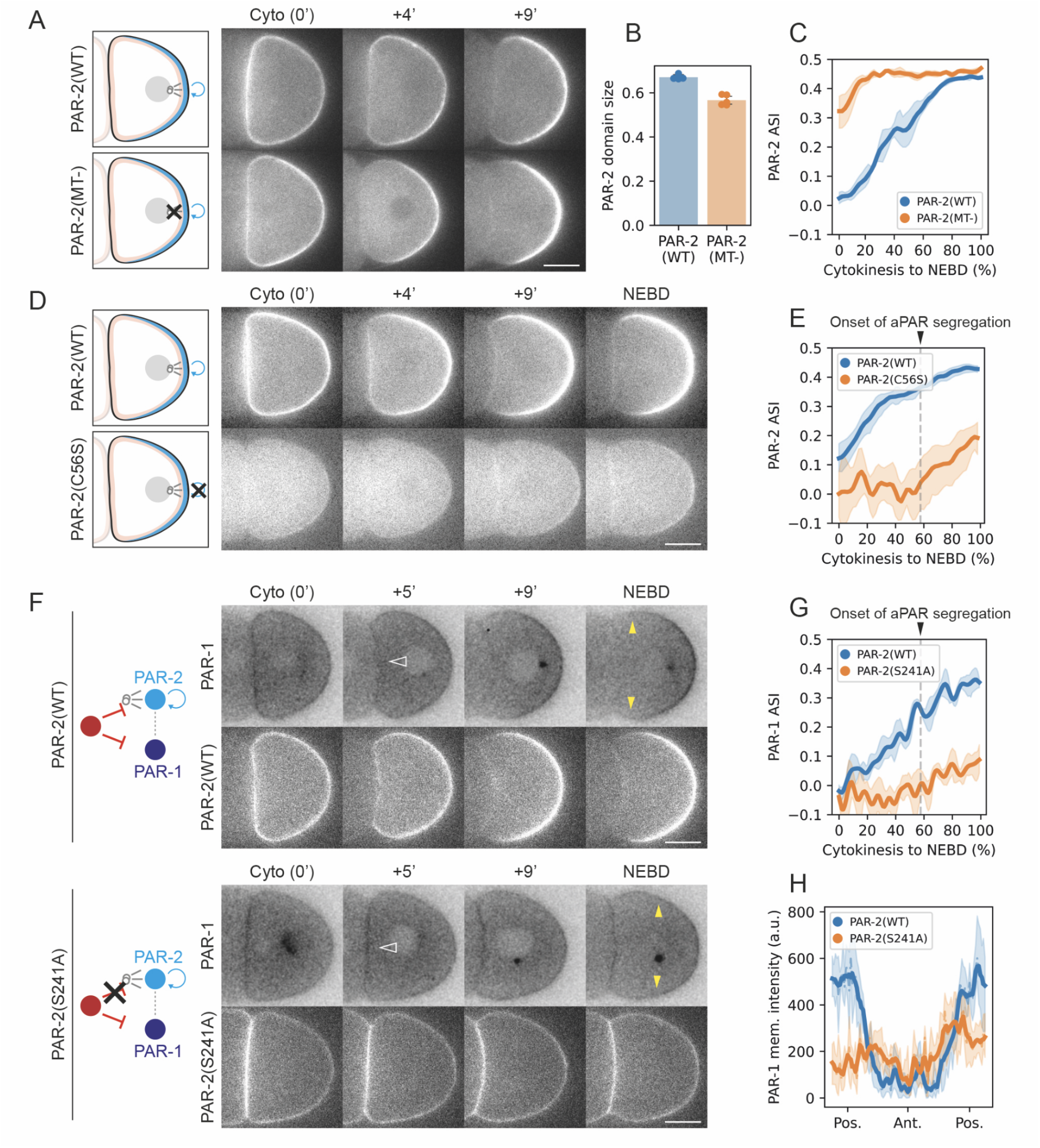
Self-organized PAR-2 domain formation is required for timely pPAR polarization. (A-C) Loss of microtubule binding activity accelerates polarization of PAR-2. A time series of midsection confocal images of embryos expressing GFP::PAR-2 (KK1273) (n=4) or GFP::PAR-2(R183-5A; MT-) (NWG0192) (n=5) (A) is shown along with quantification of PAR-2 domains size (B) and measured asymmetry (ASI = asymmetry index) over time (C). Scale bars, 10μm (D,E) Disruption of the PAR-2 RING domain blocks robust, early PAR-2 polarity. A time series of midsection confocal images of embryos expressing mNG::PAR-2 (LP637) (n=6) or mNG::PAR-2(C56S) (NWG0240) (n=8) (D) is shown along with quantification of ASI for indicated time points (E). Dotted lines indicate approximated onset of aPAR segregation as shown in Figure 1. Mean and 95% confidence interval (bootstrapped) indicated. Scale bars, 10μm. Note that the intensity range was adjusted separately for PAR-2(WT) and PAR-2(C56S) due to the relatively weaker signal of PAR-2(C56S). (F-H) Blocking PAR-2 polarization through introduction of a PKC-3 phosphorylation site mutation weakens and delays PAR-1 polarization. A time series of midsection confocal images of embryos expressing PAR-1::GFP with either mCherry::PAR-2 (NWG0332) (n=4) or mCherry::PAR-2(S241A) (NWG0344) (n=5) (F) are shown along with quantification of ASI over time (G) and PAR-1 membrane profiles at NEBD (H). Dotted lines in (G) indicate approximated onset of aPAR segregation as shown in Figure 1. Open arrowheads in *par-2(WT)* vs *par-2(S241A)* embryos highlight slower clearance of PAR-1 from the anterior (cell contact) of *par-2(S241A)* embryos. Closed yellow arrowheads roughly indicate the extent of the PAR-1 domain. A 1-pixel gaussian blur was applied and LUT inverted for PAR-1 images to improve visibility. Mean and 95% confidence interval (bootstrapped) indicated. Scale bars, 10μm

We next turned to the RING domain. We disrupted the RING domain by introducing a C56S mutation into the endogenous locus to disrupt one of the zinc-chelating cysteines (Hao et al., 2006). Importantly, polarization and asymmetric division of the zygote is largely unaffected by *par-2(C56S)* (Figure S5B) due to aPAR segregation by cortical flows (data not shown). We found that PAR-2 membrane levels decreased during cytokinesis in PAR-2(C56S) zygotes, such that by birth of P1, PAR-2(C56S) was largely cytoplasmic (Figure 3D). Thereafter, PAR-2(C56S) failed to localize to the membrane or support pPAR domain formation at the normal time. Instead, we observed weak posterior loading of PAR-2(C56S) late in the cell cycle at a time that coincided with the onset of aPAR segregation (Figure 3D, E). This suggests that, as in the zygote, PAR-2(C56S) is only able to localize to regions of the plasma membrane devoid of aPARs (Figure 3D, E) (Hao et al., 2006). Thus, we conclude that, similar to the zygote, the RING domain is required in P1 for PAR-2 to self-organize in absence of aPAR segregation, and thus required for robust and timely PAR-2 polarization in P1.

Finally, we asked how disrupting PAR-2 polarization would affect polarization of PAR-1 in P1, which as we have shown occurs with PAR-2 polarization and prior to aPAR segregation (Figure 1, Figure 3F, Figure S1E). To disrupt PAR-2 polarization, we introduced an S241A mutation into the endogenous locus, which prevents displacement from the membrane by PKC-3-dependent phosphorylation (Motegi et al., 2011). Somewhat surprisingly, despite uniform PAR-2 at the membrane, PAR-1 and PAR-6 were still able to polarize in the zygote, leading to normal asymmetric division (Figure S5C, D). We found that in the resulting *par-2(S241A)* P1 blastomeres, PAR-1 polarity was both substantially weaker and delayed such that it occurred concurrently, instead of prior to, the onset of aPAR segregation (Figure 3F-H, Figure S5F). Thus, similar to the zygote, in absence of aPAR segregation, PAR-2 self-organization promotes timely polarization of PAR-1.

### Furrow-directed flows enrich PAR-3 within the nascent cell contact

The invariable segregation of pPARs away from the nascent cell contact, which was more pronounced in *par-2(MT-)* embryos, suggested the presence of an early acting symmetry-breaking cue that biases polarization of pPARs. While imaging the segregation of aPARs at the cortex of P1 cells, we noticed that PAR-3 appeared to be transported into the nascent cleavage furrow by furrow-directed cortical flows as the zygote divided (Figure 4A, B, Movie S2), which were distinct from the “late” flows we have documented above (Figure 1F). This transport of PAR-3 was particularly prominent for PAR-3(S950A), which exhibits elevated levels at the membrane compared to wildtype (Figure 4B). Subsequent imaging of dividing embryos in cross-section revealed a marked enrichment of PAR-3 towards the leading edge of the ingressing furrow, similar to what was reported by (Pittman and Skop, 2012) (Figure 4C-E). Importantly, we found that this was not observed for the plasma membrane marker PH-PLCd1 and thus does not simply reflect an increased membrane accumulation at this site. Together, these data suggest that PAR-3 flows towards the leading edge of the cleavage furrow.

**Figure 4.**
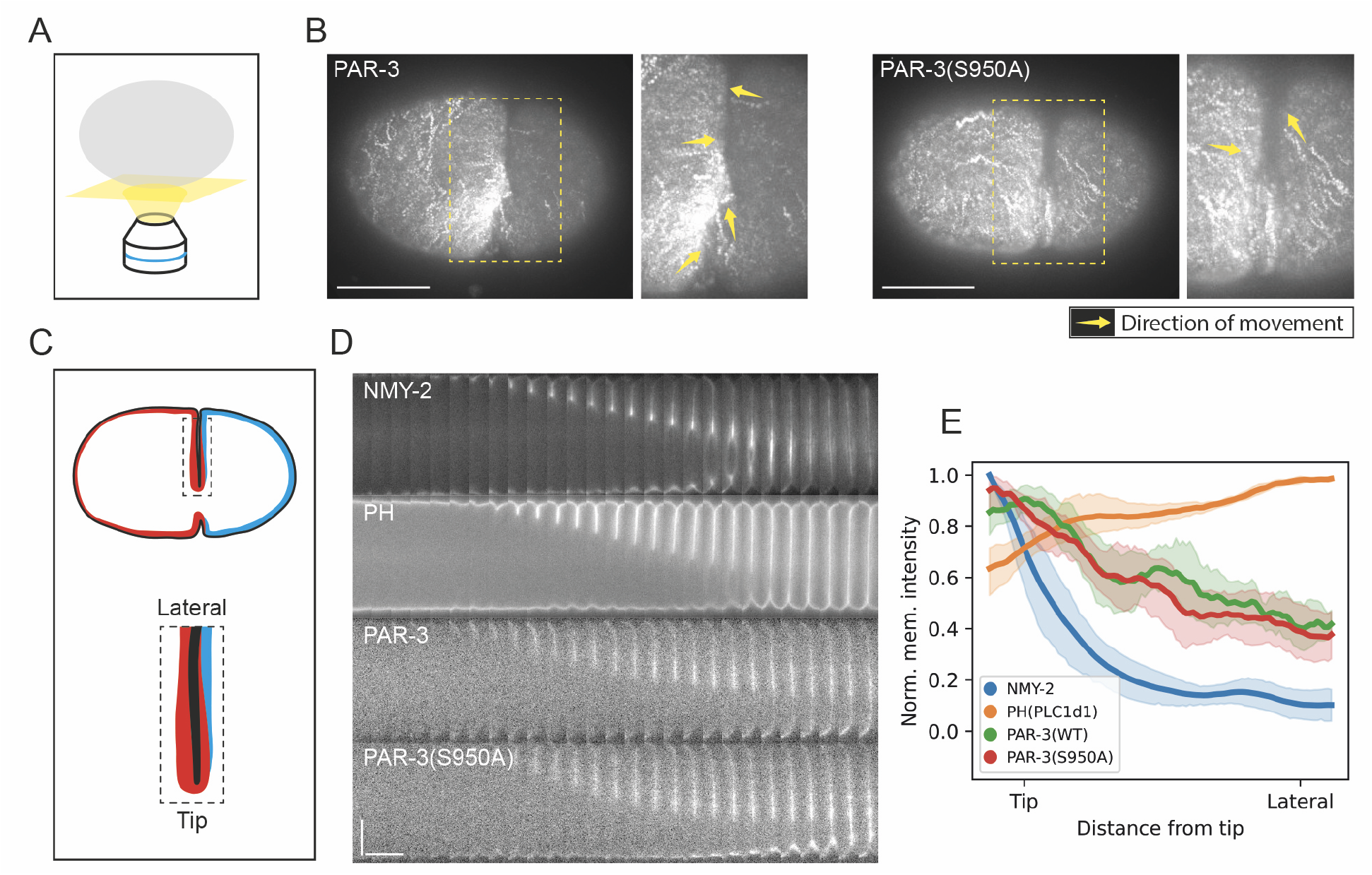
Midzone-directed cortical flows drive PAR-3 enrichment in the nascent cleavage furrow. (A) A schematic illustrating the optical section during cortical imaging. (B) Time-averaged cortical images of mNG::PAR-3 (NWG0189) (n=5) and mNG::PAR-3(S950A) (NWG0259) (n=5) spanning 182.5s during furrow closure, which reveals directed tracks of cortical clusters towards the ingression site (yellow arrows). Scale bars, 20μm. (C) A schematic illustrating the region of interest and nomenclature used when observing leading edge enrichment of molecules during furrow closure. Note that here we represent PAR-3 (red) and PAR-2 (cyan) as colocalizing on the posterior face of the furrow membranes, a point we address in Figure 5. (D) Kymograph of embryos expressing NMY-2::GFP (LP162) (n=5), GFP::PH(PLC1δ1) (OD58) (n=5), mNG::PAR-3 (NWG0189) (n=7) and mNG::PAR-3(S950A) (NWG0259) (n=6) during furrow closure. Scale bars, vertical - 10μm, horizontal 20s. Note that NMY-2 and PAR-3, but not PH, exhibit enrichment at the tip of the furrow. (E) Distribution of indicated proteins relative to the tip of the ingressing furrow as measured 30 seconds before completion of cytokinesis. Fluorescence is normalized by intensity and distance from tip to lateral edge. Mean and 95% confidence interval (bootstrapped) indicated.

We next sought to follow the fate of this accumulation in the furrow after the zygote divided. However, imaging of aPARs at the nascent cell contact of P1 is complicated by the coincidence of membranes from both AB and P1, together with much higher levels of aPARs inherited by AB. To specifically visualize proteins in P1, we devised a photobleaching technique to remove the contribution of fluorescence signals from AB (Figure 5A, Figure S6A). Briefly, just after cytokinesis, we bleached the entire AB cell, which necessarily includes a portion of the P1 membrane at the contact site. This was followed by continuous bleaching of a region of AB away from the cell contact to bleach any residual pool of fluorescent protein in AB. After a brief period (1-2 min) to allow fluorescence recovery within P1, we could then assess the distribution of protein specifically on the P1 side of the nascent contact site.

**Figure 5.**
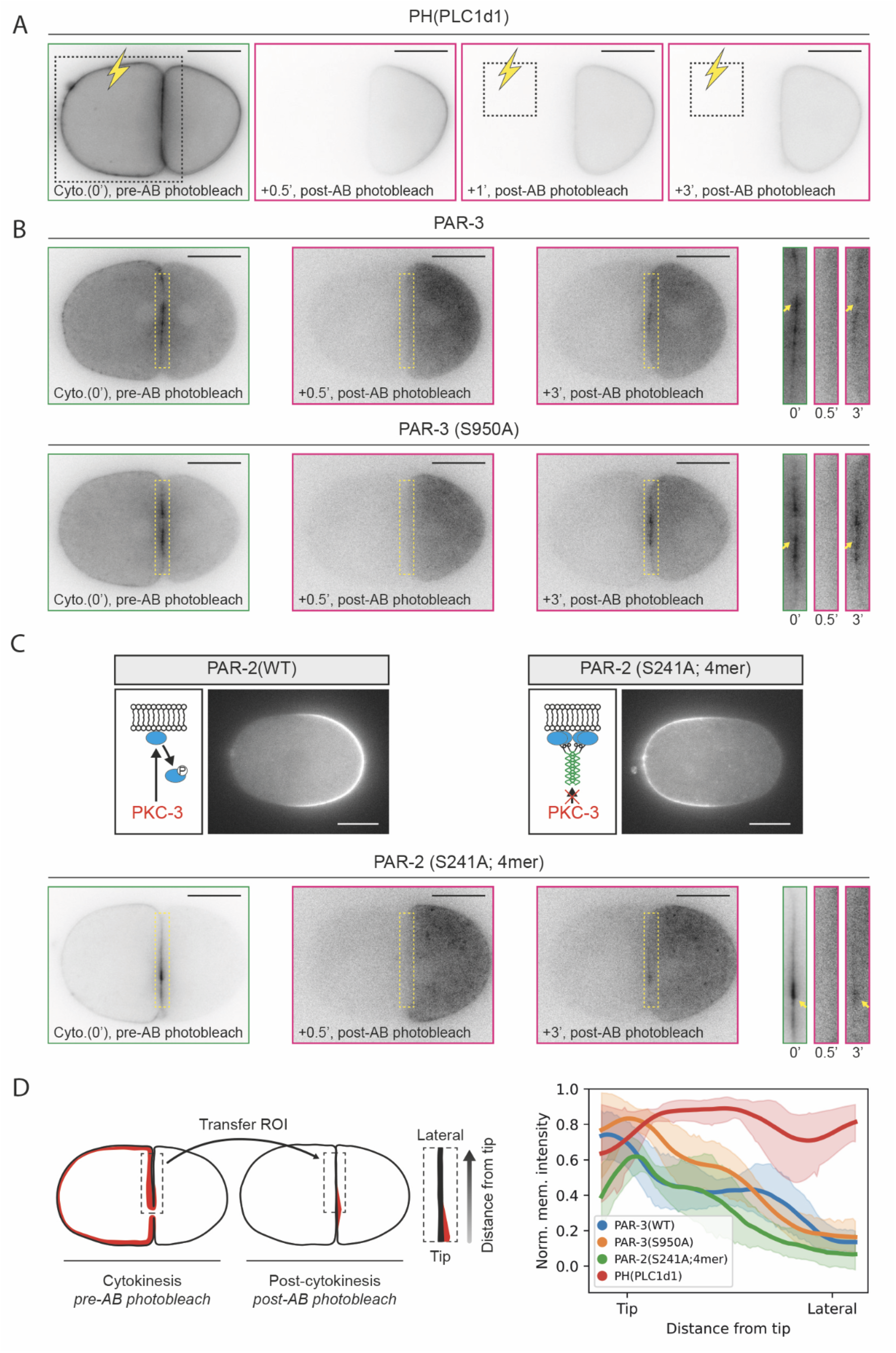
Flow-dependent accumulation establishes a persistent pool of aPARs that marks the anterior of P1. (A) Time course of photobleaching protocol used to isolate P1 membrane signal as applied to a generic plasma membrane marker, PH-PLC1δ1 (n=6). Thirty seconds after P0 cytokinesis, a region encompassing the entire AB cell, necessarily including the anteriormost region of P1 and the AB-P1 contact, was bleached (dotted box). We then allow fluorescence recovery in P1, while continually bleaching a small region of AB to further ensure that no fluorescence pool remains in AB. Consequently, any remaining signal at the cell contact should reflect proteins from P1. Note the excess fluorescence at the cell contact due to the presence of two abutted membranes is reduced and the PH-PLC1δ1 signal becomes uniform in P1. See also Figure S6. (B) Representative example of pre- (0’) and post-recovery (0.5’, 3’) images of embryos expressing mNG::PAR-3 (NWG0189) (n=8) and mNG::PAR-3(S950A) (NWG0259) (n=6). Insets highlight accumulation of signal towards the furrow closure site at the AB-P1 contact. Yellow arrows indicate where molecules accumulate during furrow closure leads directly to a corresponding enrichment in P1 post-cytokinesis, as revealed by the photobleaching protocol. See also Figure S6. (C) (Top) A schematic and midsection confocal image showing localizations of mNG::PAR-2 (LP637) and mNG::PAR-2 (S241A; 4mer) (NWG0473) in the zygote, which localize to the posterior and anterior respectively. (Bottom) Representative examples of pre- (0’) and post-recovery (0.5’, 3’) images of embryos expressing mNG::PAR-2 (S241A; 4mer) (NWG0473) (n=7) as in (B). (D) Fluorescence profiles of PAR-3, PAR-3(S950A), and PAR-2(S241A; 4mer), are persistently enriched at the site of furrow closure, but not PH-PLC1δ1, following the photobleaching protocol shown in (A). (Left) A schematic of the quantification protocol is shown. Briefly, an ROI was specified as indicated at the time of furrow closure and applied to the post-recovery image frame (3’-5’) to obtain a profile of fluorescence signal relative to the point of furrow closure. (Right) Quantification of membrane profiles from embryos corresponding to conditions shown in (A-C). Profiles were normalized by intensity and distance. Mean and 95% confidence interval (bootstrapped) indicated. Scale bars, 20μm

Applying this approach to PH-PLCd1 revealed a uniform enrichment throughout the plasma membrane of P1 (Figure 5A, Figure S6, Movie S3). By contrast, we found a persistent pool of PAR-3, and to a lesser extent PAR-6, localized to the contact site (Figure 5B, Figure S6B, C, Movie S3). Both visual inspection and quantification of spatial intensity profiles along the contact site revealed that the peak of PAR-3 accumulation was generally biased towards the site of furrow closure, which again was not observed for a generic plasma membrane marker (Figure 5A, B, D, Figure S6). Taken together, our data suggest that cortical flows into the nascent cleavage furrow and the resulting enrichment of PAR-3 on furrow membranes ultimately leads to the formation of a pool of PAR-3 that marks the anterior of the newly born P1 cell.

To confirm that furrow-directed transport was sufficient to account for this enrichment of PAR-3 at the nascent cell contact, we asked whether we could induce enrichment of a molecule at the contact site simply by allowing it to “sense” flows. To this end, we used an engineered form of PAR-2, PAR-2(S241A; 4mer) which undergoes highly efficient transport by cortical flow and is insensitive to displacement from the membrane by PKC-3 phosphorylation (Illukkumbura et al., 2022). Consequently, unlike PAR-2(WT), PAR-2(S241A; 4mer) was efficiently segregated by cortical flows into the anterior of the zygote along with aPARs (Figure 5C). We found that in P1, PAR-2(S241A; 4mer) exhibited a near identical pattern of accumulation towards the nascent cell contact as PAR-3 (Figure 5, Figure S6), confirming that advection of molecules by cortical flows is sufficient to drive accumulation at the nascent cell contact.

Taken together, our data suggests that advection of PAR-3 by cleavage furrow-directed cortical flows establishes a locally enriched and intrinsically stable pool of PAR-3 at the nascent cell contact. This pool persists after furrow closure and is thereby positioned to mark the anterior (nascent cell contact) of P1, and is well placed to provide directional cues to bias downstream polarization pathways.

### Accumulation of aPARs at the cleavage furrow can induce mirror-symmetric polarity in symmetrically dividing cells

Cytokinesis has been implicated as a polarity cue in a variety of systems, including neuronal polarization in *Drosophila*, lumen formation and yeast bud scars (Etienne-Manneville, 2004; Harris and Tepass, 2010; Li et al., 2014; Luján et al., 2016; Meitinger et al., 2014; Miller et al., 2020; Pollarolo et al., 2011; Rathbun et al., 2020; Schlüter et al., 2009; Wang et al., 2014). One striking example is the so-called mirror symmetric division, in which two daughter cells polarize in opposing directions relative to the nascent cell contact (Buckley et al., 2013; Tawk et al., 2007). Roles for the spindle midzone, midbody remnants and cell contact have been reported (Buckley et al., 2013; Li et al., 2014; Liang et al., 2022; Luján et al., 2016; Schlüter et al., 2009; Wang et al., 2014), which likely play complementary roles during establishment of cell and tissue polarity (Buckley and St Johnston, 2022). However, whether advection of polarity proteins by furrow-directed flows also plays a role has not been explicitly tested.

We wondered whether furrow-directed flows could, at least in principle, be sufficient to induce mirror-symmetric polarization. To this end, we suppressed polarity and asymmetric division in the zygote using the *pkc-3*^*AS*^ allele to generate symmetric daughter pairs (Figure 6A) (Ng et al., 2022). In these symmetric cell pairs, we observed aggregates of PAR-3 molecules that clearly moved inwards with the ingressing furrow membranes during cytokinesis (Figure 6B, Movie S4), which led to accumulation at the leading edge of the cleavage furrow and ultimately enrichment of PAR-3 at the nascent cell contact (Figure 6, Figure S7). When 1NA-PP1 was washed out to reactivate PKC-3, the cell pairs invariably developed mirror-symmetric polarity, with PAR-2 segregating away from the nascent division site (Figure 6, Figure S7, Movie S4). Consequently, neither fate of the daughter cells nor pre-existing polarity inherited from polarization of the zygote is required for PAR polarization with respect to the furrow, consistent with de novo polarization by the cleavage furrow.

**Figure 6.**
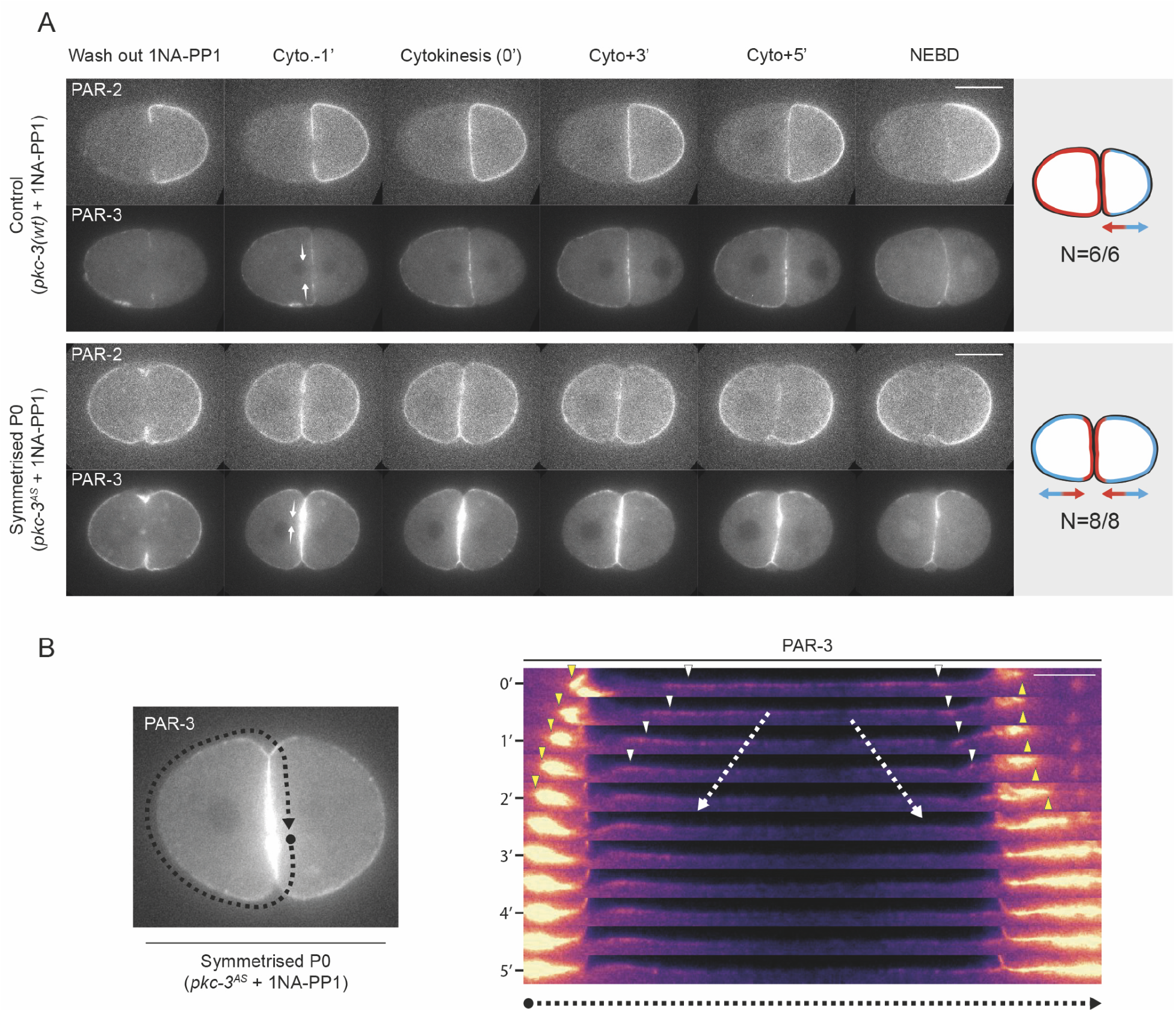
Advection of PAR-3 by furrow-directed flows can induce mirror-symmetric polarization in daughter cells. (A) Time series of midsection confocal images of embryos at indicated time points expressing mCherry::PAR-2 and mNG::PAR-3 in either PKC-3(WT) (NWG0453) or PKC-3^AS^ (NWG0458) backgrounds, when the zygote is treated with 100μM 1NA-PP1 and washed out during furrow formation. White arrows indicate the direction of furrow closure, which is where PAR-3 is enriched. Schematics of the resulting polarity shown at right, graded arrows indicate orientation of PAR polarity, aPARs are labeled as red and pPARs as cyan. Scale bars, 20μm (B) Kymograph of PAR-3 distributions for the indicated ROI (dotted line with arrowhead) during cytokinesis following washout of 1NA-PP1 in a symmetrized PKC-3^AS^ embryo (the same embryo shown in (A)). Note that PAR-3 flows towards the leading edge of the forming furrow (white arrowheads), leading to clearance of PAR-3 from the pole and accumulation of PAR-3 at the leading edge (yellow arrowheads). Scale bar, 10μm

Taken together, our data suggest that advection of polarity molecules by furrow-directed cortical flows is sufficient to induce mirror symmetric polarity in daughter cells and thus could be a broadly conserved mechanism for inducing and/or re-orienting molecular asymmetries during cell division.

## Discussion

Here, we present the first comprehensive description of the process of polarization in the germline blastomere P1 (Figure 7), which proceeds via the following steps: first, advection of PAR-3 by “early” cleavage furrow-directed cortical flows drives accumulation of a pool of PAR-3 and associated aPARs at the nascent cell contact, marking the putative anterior. Outside of this selective enrichment, aPAR membrane concentrations are initially low throughout the rest of the cell, while pPARs are uniformly high (Figure 7). Next, PAR-2 self-organization drives timely formation of a posterior pPAR domain, the position of which is consistent with being biased by aPARs at the cell contact. Concurrently, aPARs load progressively throughout the plasma membrane of P1, leading to overlap between aPAR and pPAR proteins at the posterior. Finally, pPAR-dependent exclusion of aPARs and “late” cortical flows combine to drive bulk segregation of aPARs into a visible anterior domain, resulting in the characteristic mutually exclusive distributions of aPARs and pPARs. Although technical limitations prevented more extensive analysis in P2, we were able to obtain data which suggests that this basic picture is preserved (Figure S8). Notably, in addition to polarity being sensitive to disruption of cortical flows and aPAR exclusion (Figure S8B), we also found that PAR-2 was initially biased away from the contact site as in P1, before eventually reversing due to the signaling from its neighboring endoderm precursor EMS (Arata et al., 2010), suggesting that the furrow-associated cue is also active but ultimately overridden in P2 (Figure S8C).

**Figure 7.**
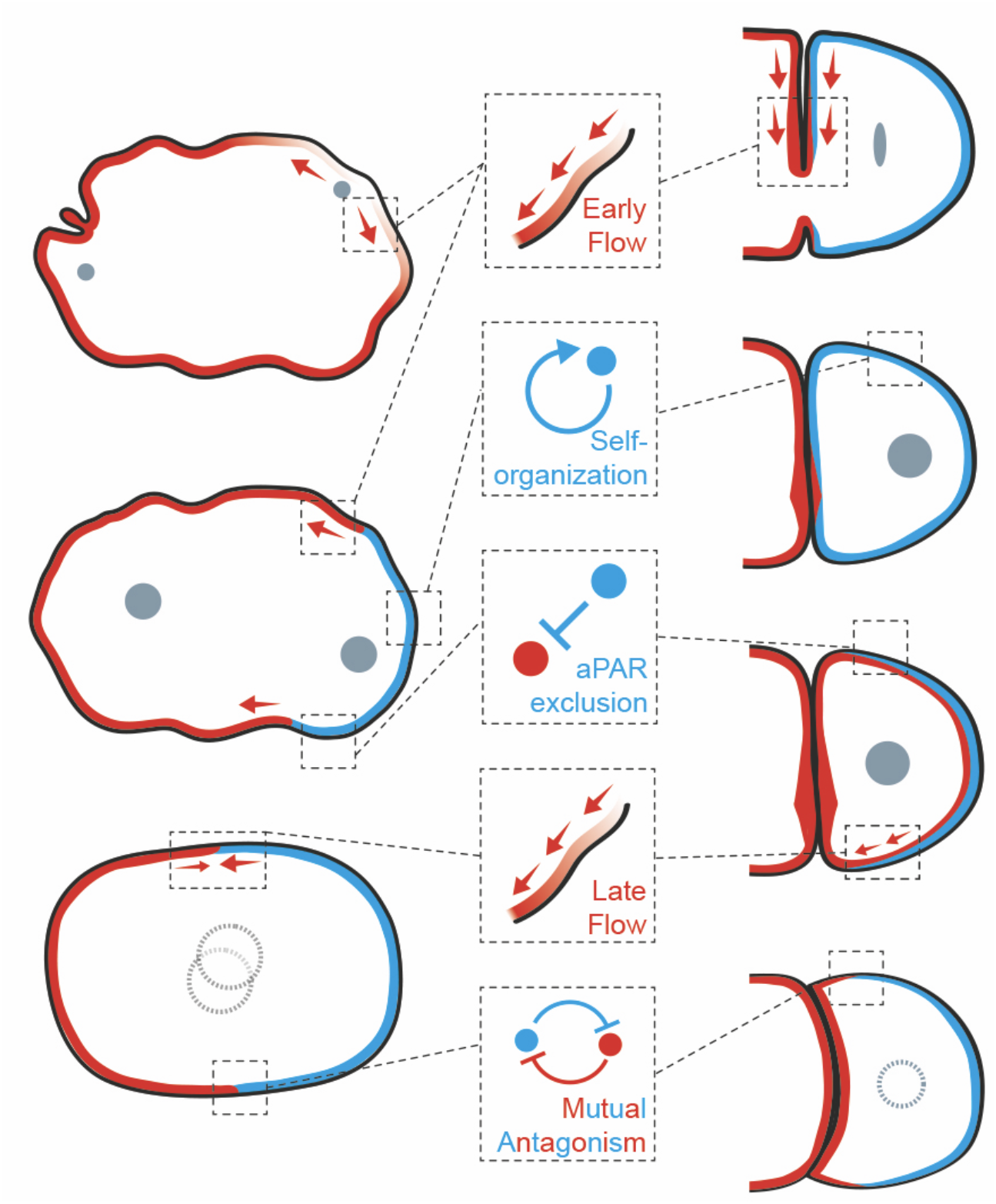
Comparison of PAR polarization in P lineage blastomeres. Schematic comparing and depicting the working model for PAR polarization in P lineage blastomeres. (Left) In the zygote, prior to symmetry breaking (SB), aPARs are initially enriched on the cortex. SB is triggered by centrosomes, which induces “early” cortical flows that segregate aPARs to the anterior. Self-organization of PAR-2 at the posterior also helps form and stabilize a posterior pPAR domain, which excludes aPARs from the posterior. By mitosis, aPARs and pPARs display mutually exclusive localizations, which are maintained via cross-inhibition. The boundaries of each PAR domain are also stabilized by “late” cortical flows which are directed towards the midline. (Right) During P0 cytokinesis, “early” cleavage furrow-directed flows lead to accumulation of PAR-3 at the leading edge, which concentrates PAR-3 at the nascent cell contact (anterior) of P1. By completion of P0 cytokinesis, these “early” flows cease, leading to low aPAR levels across the membrane of the cell apart from enrichment at the cell contact. In contrast, pPARs are uniformly distributed throughout P1 initially. Soon after, pPARs polarize away from the cell contact (anterior), matching where aPARs accumulate during “early” flows, and are dependent on PAR-2 self-organization for timely formation of a pPAR domain. Concurrently, aPARs load throughout the cell, resulting in overlap of aPARs and pPARs at the posterior. Finally, “late” flows and posterior exclusion of aPARs by pPARs segregates aPARs towards the anterior, resulting in the stereotyped mutually-exclusive localization of aPARs and pPARs.

### Diverse patterns of PAR polarization arise from a common set of core mechanisms

Compared to the zygote, the pattern of polarization in P lineage blastomeres differs in several notable ways (Figure 7). First, the initial state of PAR proteins at the membrane prior to polarization is inverted relative to the zygote, i.e. with pPARs high and aPARs low. Second, in contrast to the zygote in which aPAR and pPAR domains emerge together as aPARs are segregated into the anterior by cortical flows, emergence of aPAR and pPAR domains in P lineage blastomeres is decoupled. Third, there are extensive periods in which aPAR and pPAR localisations are not mutually exclusive. Finally, rather than being linked to a single period of polarizing cortical flow, polarity in P1 is linked to two distinct periods of cortical flow separated by a temporal gap: an “early” cleavage furrow-directed cortical flow breaks symmetry by driving an accumulation of a PAR-3 pool at the cell anterior, while a second period of “late” flow helps drive bulk segregation and coalescence of aPARs into an anterior domain. Zygotes also exhibit so-called late flows. However, unlike those found in the P lineage blastomeres, late flows in the zygote closely follow the cessation of early flows and are oriented bidirectionally from both the anterior and posterior towards the embryo midline, where they are primarily thought to reenforce and stabilize the PAR boundary (Sailer et al., 2015).

Given such differences, it is rather striking that the core mechanisms of PAR protein segregation are largely conserved between these cells (Figure 7): (1) Polarization begins with an initial symmetry breaking event involving segregation of aPARs by cortical flows; (2) Polarity is promoted by PAR-2 self-organization; (3) Segregation of aPARs is driven by semi-redundant contributions of cortical flow and exclusion by pPARs.

### Patterns of polarity emergence are defined by the cell state

We suggest that the disparate manifestations of polarity establishment in these cells arise not from distinct polarity mechanisms, but from distinct peculiarities of each cell’s life history. For example, in the zygote, the earlier timing of aPAR domain formation can be explained by the fact that aPAR membrane levels are already high at the onset of cortical flows, which allows them to be segregated into an aPAR domain early in the cell cycle. In contrast, in later P lineage blastomeres, the cessation of furrow-associated flows occurs prior to bulk loading of aPARs onto the plasma membrane. Thus, bulk aPAR segregation is delayed until the onset of late flows. Consistent with this interpretation, disruption of “early” flows in the zygote shifts polarization to a P1-like program: PAR-2 now polarizes prior to aPAR segregation, leading to transient overlap of aPARs and pPARs in the posterior, until the combined action of antagonistic pPAR activity and a period of “late” cortical flows drive aPAR exclusion from the posterior (Zonies et al., 2010).

Similarly, the distinct initial states of the zygote vs P lineage blastomeres could explain the rather striking difference in behavior of PAR-2(MT-) mutants: whereas the protection against PKC-3 provided by MT association is an advantage in the zygote, in which PAR-2 must invade an aPAR-dominated cortex, it may be a disadvantage in P1, in which aPAR levels on the membrane are low initially and PAR-2 must be cleared from the anterior. Consistent with this interpretation, we have found that early polarization of P1 in PAR-2(MT-) mutants can be suppressed by mild aPAR depletion (data not shown).

Finally, our data suggest that the mutually exclusive nature of aPAR and pPAR domains, enforced through mutual antagonism, is a tunable parameter of the system rather than an intrinsic requirement for the establishment of PAR domains per se. Notably, we found that in P1, formation of aPAR and pPAR domains is temporally uncoupled, which leads to overlap between the two domains for much of the cell cycle. The canonical mutually exclusive localizations of aPARs and pPARs, which is more typical of the zygote, only appears late in P1. Intriguingly, in the zygote, such uncoupling has been observed when anterior and posterior PAR proteins are out of balance. In these cases, aPARs and pPARs initially overlap but are sometimes able to resolve into mutually-exclusive domains later in the cell cycle (Dickinson et al., 2017; Lim et al., 2021), which is almost identical to what we observe in wild-type P1 cells. Therefore, the temporal progression towards an increasing mutual exclusion of aPARs and pPARs that we observe in P1 could suggest that embryos temporally optimize levels of PAR cross-talk, potentially adding to already established roles for cell cycle-dependent regulation of PAR polarity and cell fate specification (Bell et al., 2015; Carvalho et al., 2015; Dickinson et al., 2017; Klinkert et al., 2019; Peglion and Goehring, 2019; Wirtz-Peitz et al., 2008).

### Furrow-induced flows as a paradigm for division-linked symmetry breaking

Work in many systems has demonstrated the coupling of cell polarity to cell division, which presumably allows coordination of cellular geometry in tissues that are actively dividing (Buckley and St Johnston, 2022). This division-linked polarity has been linked to a variety of spatial cues, including cytokinesis remnants, the spindle midzone, the midbody, and/or contact-dependent signaling between newly generated daughters (Etienne-Manneville, 2004; Harris and Tepass, 2010; Li et al., 2014; Luján et al., 2016; Meitinger et al., 2014; Miller et al., 2020; Pollarolo et al., 2011; Rathbun et al., 2020; Schlüter et al., 2009; Wang et al., 2014), which likely act together to ensure proper cell polarization (Buckley and St Johnston, 2022). In many of these cases polarization has been linked to the accumulation of polarity related molecules, including PAR-3, within the cleavage furrow and/or nascent contact site (Buckley et al., 2013; Harris and Peifer, 2004; Liang et al., 2022; Tawk et al., 2007). Our data further suggests that advection of polarity proteins by furrow-directed cortical flow also may play a role, by providing a simple mechanism to concentrate such molecules within the furrow to ensure tight coupling between division and the polarity axis.

While it is difficult to fully exclude other potential mechanisms of PAR-3 accumulation at the nascent cell contact in P1, several lines of evidence favor our model of flow-dependent accumulation. Most notably, PAR-3 visibly flows into the cleavage furrow and accumulates towards the leading edge during cytokinesis, which directly leads to a corresponding enrichment at the P1 cell anterior following cell division. In addition, although there is evidence for contact-dependent signaling between somatic and P lineage blastomeres (Anderson et al., 2008; Arata et al., 2010; Klompstra et al., 2015), these pathways generally inhibit rather than enhance aPAR accumulation at cell contacts. We would point out that when isolated P lineage blastomeres are placed in contact with non-E lineage cells, they polarize in random directions with respect to the cell contact, suggesting a lack of contact-dependent signaling in such contexts (Arata et al., 2010).

Given the ubiquity of furrow-directed flows in dividing cells (Bray and White, 1988; Cao and Wang, 1990; Chen et al., 2008; Yumura, 2001), it would be surprising that they are not used as a general mechanism for spatial patterning during development, either to concentrate molecules that may help promote cytokinesis (Khaliullin et al., 2018; Longhini and Glotzer, 2022), or as seen here to induce asymmetry in daughter cells.

## Conclusion

In summary, through comparative analysis of polarization in the *C. elegans* germ lineage, we have identified conserved mechanisms of polarization, revealed how they are adapted to new contexts, and identified a role for furrow-directed cortical flows during cytokinesis in defining cell division-linked cellular patterning.

## Methods

### *C. elegans* strains and culture conditions

*C. elegans* strains were maintained on OP50 bacterial lawns seeded on nematode growth media (NGM) at 20°C or 15°C (for experiments involving temperature sensitive mutants) under standard laboratory conditions (Stiernagle, 1999). OP50 bacteria were obtained from CGC. Zygotes were obtained from hermaphrodites unless otherwise noted. Analysis of embryos precludes determination of animal sex. All strains used in this study are indicated in Table S1.

### Strain construction

Mutation by CRISPR-Cas9 was performed based on the protocol published by (Arribere et al., 2014). Briefly, tracrRNA (IDT DNA, 0.5 μL at 100 μM) and crRNA(s) for the target (IDT DNA, 2.7 μL at 100μM) with duplex buffer (IDT DNA, 2.8μL) were annealed together (5 min, 95°C) and then stored at room temperature until required. An injection mix containing Cas9 (IDT DNA, 0.5μL at 10mg/mL), annealed crRNA, tracrRNA, and the repair template (IDT Ultramer) was incubated at 37°C for 15 min and centrifuged to remove debris (10 min, 13,000 rpm). Young gravid adults were injected along with either a *dpy-10* or *unc-58* co-CRISPR injection marker (Arribere et al., 2014) and mutants identified by PCR and sequence verified.

### RNAi culture conditions

RNAi by feeding was performed according to previously described methods (Kamath and Ahringer, 2003). Briefly, HT115(DE3) bacterial feeding clones were inoculated from LB agar plates to LB liquid cultures and grown overnight at 37ºC in the presence of 50 μg/mL ampicillin (until a fairly turbid culture is obtained). To induce high dsRNA expression, bacterial cultures were then treated with 1 mM IPTG before spotting 150μL of culture onto 60 mm NGM agar plates (supplemented with 10 μg/ml carbenicillin, 1 mM IPTG) and incubated for 24 hr at 20ºC. L3/L4 larvae were then added to RNAi feeding plates and incubated for 24-32 hours at either 20ºC or 25ºC.

### Dissection and mounting for microscopy

Embryos were obtained by dissecting adult worms in 8-10μL of egg buffer (118 mM NaCl, 48 mM KCl, 2 mM CaCl2 2 mM MgCl2, 25 mM HEPES, pH 7.3), and mounted with 18.8 μm (cortex imaging) or 20 μm (midplane imaging) polystyrene beads (Polysciences, Inc.) between a slide and coverslip as in (Rodriguez et al., 2017), and sealed using VALAP (1:1:1, vaseline:lanolin:paraffin wax).

For laser ablation mediated extrusion of ABa and ABp in 4-cell stage embryos, embryos were dissected in 8-10 μL of Shelton’s Growth Medium (Edgar and Goldstein, 2012) and mounted with 20 μm polystyrene beads (Polysciences, Inc.) between a slide and coverslip, and sealed using VALAP as above.

For acute drug treatment and washout experiments, embryos were first permeabilized using either *ptr-2* or *perm-1* fRNAi. Embryos were then dissected in 8-10 μL of Shelton’s Growth Medium (with or without drugs) (Edgar and Goldstein, 2012), and mounted with 18.8 μm polystyrene beads between a large and small coverslip sealed on two parallel edges with VALAP as in (Goehring et al., 2011a). Buffer exchange is achieved through capillary action by placing a drop of solution at one side of the sample, and touching a piece of filter paper at the opposite side.

Type of drug used, the respective concentration and the relative timings for drug addition/ washout for each experiment are as follows. For inhibiting PAR-2 polarization in *pkc-3*^*AS*^ P1 embryos, 20 μM of 1NA-PP1 (Calbiochem, 529579) was washed in soon before P0 cytokinesis completion. To depolymerize microtubules in P1, 10 μg/ml nocodazole was washed in soon before P0 cytokinesis completion. To reactivate PAR polarity in symmetrized 2-cell *pkc-3*^*AS*^ embryos, worms were dissected directly in 100 μM of 1NA-PP1 (Calbiochem, 529579), zygotes were then tracked and soon after cytokinesis onset, the drug was washed out.

### Temperature upshift for temperature sensitive alleles

Rapid temperature upshift for *nmy-2(ts)* alleles was achieved by first preheating one objective lens (100X) to 25.5ºC, and controlling the room temperature at 18.5ºC. Following, embryos were dissected and mounted onto an objective lens without a temperature collar (∼18.5ºC), and zygotes were tracked and imaged through cytokinesis. 3 minutes after cytokinesis completion in P1 (and 5 minutes for P2), the objective lens was swapped with the preheated one (which roughly takes 30 to 60s), and imaging was continued. The 3 (or 5) minute buffer time was to ensure that cells do not refuse following cytokinesis as previously reported (Liu et al., 2010). Disruption of *nmy-2(ts)* activity was confirmed by scoring cytokinesis failure following upshift.

### Microscopy - live imaging

Midsection confocal images were captured on a Nikon TiE with a 100x/1.40 NA oil objective, further equipped with a custom X-Light V1 spinning disk system (CrestOptics, Rome, Italy) with 50 μm slits, Obis 488/561 fiber-coupled diode lasers (Coherent, Santa Clara, CA) and an Evolve Delta EMCCD camera (Photometrics, Tuscon, AZ). Imaging systems were run using Metamorph (Molecular Devices, San Jose, CA) and configured by Cairn Research (Kent, UK). Filter sets were from Chroma (Bellows Falls, VT): ZT488/561rpc, ZET405/488/561/640X, ET535/50m, ET630/75m. Imaging of cortical flows using NMY-2::GFP embryos was achieved as above, but by acquiring a stack instead with 5s intervals. The maximum intensity projection of the stack for each time point was then used for further analysis (Pimpale et al., 2020). All embryos were imaged with a 20ºC temperature collar, except for the temperature sensitive experiments, to which the temperature collar was set at 25.5ºC.

Cortical imaging was carried out with a 100x 1.49 NA TIRF objective on a Nikon TiE microscope equipped with an iLas2 TIRF unit (Roper), a custom-made field stop, 488 or 561 fiber coupled diode lasers (Obis), and an Evolve 512 Delta EMCCD camera (Photometrics), controlled by Metamorph software (Molecular Devices) and configured by Cairn Research. Filter sets were from Chroma: ZT488/561rpc, ZET488/561x, ZET488/561m, ET525/50m, ET630/75m, ET655LP. Images were captured in bright field, GFP/mNG (ex488/ZET488/561m), RFP/mKate/mCherry (ex561/ZET488/561m).

### Microscopy - cell specific photobleaching

Contribution of fluorescence signal from AB (2-cell stage) or EMS (4-cell stage) to the AB-P1 or EMS-P2 contact site was removed using photobleaching. First, a bleaching box was specified which encompassed AB and the AB-P1 contact site, or EMS and the EMS-P2 contact site, and was subsequently subjected to photobleaching using a 473 nm laser at full power for 10 repetitions to ensure near complete depletion of signal. Following, a new smaller bleach box was set, replacing the old one, which encompasses a small region on the anterior end of AB or EMS, and was continually bleached every 30s between imaging. This was to continually deplete the fluorescence pool in AB or EMS, which could contribute to fluorescence recovery at the contact site.

### Microscopy - laser ablation mediated extrusion of ABa and ABp

To extrude ABa and ABp, we ablated a small circular region encompassing the anterior side of ABa with the eggshell, and the dorsal side of ABp with the eggshell with a 355nm laser using a iLas2 Pulse targeted illumination system (Roper). This causes the cells to lyse and to extrude itself out of the opening generated when ablating the eggshell.

### Image analysis - particle image velocimetry

The mean flow velocity of NMY-2 was calculated through a particle image velocimetry (PIV) analysis using the PIVlab MATLAB plugin (Thielicke and Stamhuis, 2014). Briefly, maximum intensity projection of stacks was first processed using background subtraction and a 1 pixel median filter in Fiji. A box encompassing the entirety of P1 was selected as the ROI and was subjected to a high-pass filter with other preprocessing filters disabled. A FFT phase-space PIV algorithm was utilized with a 3 step multi pass linear window deformation, with a final area of 16 pixels. A velocity filter with a limit of 10 pixels per frame was applied together with a standard deviation filter, where any vectors that are 3 standard deviations from the mean are removed. Interpolation was used to replace lost vectors.

### Image analysis - quantification of membrane profile

Raw or SAIBR processed images were used for quantification (Rodrigues et al., 2022). In order to measure cortical concentrations, a 100-pixel-wide (15.5 μm) line following the membrane around the embryo was computationally straightened, and a 20-pixel-wide (3.1 μm) rolling average filter was applied to the straightened image. Intensity profiles perpendicular to the membrane at each position were fit to the sum of a Gaussian component, representing membrane signal, and an error function component, representing cytoplasmic signal, and a constant, representing background signal. Membrane concentrations at each position were calculated as the amplitude of the Gaussian component. This protocol is similar to previously published methods (Gross et al., 2019; Reich et al., 2019) and identical to (Ng et al., 2022).

### Image analysis – defining anterior and posterior poles in the zygote

The overall geometry of the zygote was first defined by fitting the shape of the ROI to an ellipsoid. The anterior and posterior pole was defined as the ROI coordinate closest to the tip of each side of the major axis.

### Image analysis – alignment of time series data in P1

Because PAR domains are more variable in position during polarization in P1, membrane profiles were aligned throughout the cell cycle for each embryo, followed by alignment between embryos to ensure accurate representation of PAR polarization dynamics. To align membrane concentration profiles throughout the cell cycle of an embryo, membrane profiles that were adjacent in time were averaged; individual profiles within that time span were aligned to the mean, and this process was iterated until a lowest mean squared error was obtained. To align membrane profiles between embryos, an averaged membrane profile for time points around NEBD was used as a reference for each embryo, and aligned in a way identical to before, i.e. the membrane profiles of individual embryos were aligned to the mean of all embryos, and the process was iterated until the lowest mean squared error was achieved. Membrane profiles were also geometrically corrected, through automated tracing of the ROIs in both clockwise and anti-clockwise fashion, and to flip membrane profiles of individual embryos so that the average of all embryos has the lowest mean squared error when aligned.

### Image analysis – ASI

For calculating ASI, the following equation was used: ASI = (A – P) / (2 * (A + P)), where A and P denote the sum of fluorescence signals in the anterior or posterior 30% of the cell respectively (Rodriguez et al., 2017).

As embryos vary slightly in their cell cycle length, to match ASI from cytokinesis to NEBD between different embryos, a cubic interpolation was used to normalize data to the same length.

### Image analysis - domain size

For calculating domain size, fluorescence profiles were first fitted to an error function with a defined gradient center and width. The domain size was calculated as total length of *fluorescence profile – (gradient center – 0*.*75 * gradient width)*. Domain size is then defined as a fraction of the embryo perimeter (Ng et al., 2022; Rodriguez et al., 2017).

## Contributions

Conceptualization: K.N., N.W.G.; Methodology: K.N.; Formal analysis: K.N.; Investigation: K.N.; Resources: K.N., N.H., T.B., J.B.-P.; Writing – original draft preparation: K.N., N.W.G.; Writing – review and editing: K.N., N.W.G.; Supervision: N.W.G.; Project administration: N.W.G.; Funding acquisition: N.W.G.

## Acknowledgements

We thank Nic Tapon and Clare Buckley for comments on the manuscript and Lesilee Rose for discussions of unpublished data. Some strains were provided by the Caenorhabditis Genome Center (CGC), which is funded by NIH Office of Research Infrastructure Programs (P40 OD010440). This work was supported by the Francis Crick Institute (N.W.G.), which receives its core funding from Cancer Research UK (CC2119), the UK Medical Research Council (CC2119), and the Wellcome Trust (CC2119). For the purpose of Open Access, the author has applied a CC BY public copyright license to any Author Accepted Manuscript version arising from this submission.

## Data Availability

Source code and documentation will be made available on GitHub.

## References

Anderson, D.C., Gill, J.S., Cinalli, R.M., Nance, J., 2008. Polarization of the C. elegans embryo by RhoGAP-mediated exclusion of PAR-6 from cell contacts. Science 320, 1771–1774.

Arata, Y., Hiroshima, M., Pack, C.-G., Ramanujam, R., Motegi, F., Nakazato, K., Shindo, Y., Wiseman, P.W., Sawa, H., Kobayashi, T.J., 2016. Cortical polarity of the RING protein PAR-2 is maintained by exchange rate kinetics at the cortical-cytoplasmic boundary. Cell Rep. 16, 2156–2168.

Arata, Y., Lee, J.-Y., Goldstein, B., Sawa, H., 2010. Extracellular control of PAR protein localization during asymmetric cell division in the C. elegans embryo. Development 137, 3337–3345.

Arribere, J.A., Bell, R.T., Fu, B.X., Artiles, K.L., Hartman, P.S., Fire, A.Z., 2014. Efficient marker-free recovery of custom genetic modifications with CRISPR/Cas9 in Caenorhabditis elegans. Genetics 198, 837–846.

Beatty, A., Morton, D., Kemphues, K., 2010. The C. elegans homolog of Drosophila Lethal giant larvae functions redundantly with PAR-2 to maintain polarity in the early embryo. Development 137, 3995–4004.

Bell, G.P., Fletcher, G.C., Brain, R., Thompson, B.J., 2015. Aurora kinases phosphorylate Lgl to induce mitotic spindle orientation in Drosophila epithelia. Curr. Biol. 25, 61–68.

Boyd, L., Guo, S., Levitan, D., Stinchcomb, D.T., Kemphues, K.J., 1996. PAR-2 is asymmetrically distributed and promotes association of P granules and PAR-1 with the cortex in C. elegans embryos. Development 122, 3075–3084.

Bray, D., White, J.G., 1988. Cortical flow in animal cells. Science 239, 883–888.

Buckley, C.E., Ren, X., Ward, L.C., Girdler, G.C., Araya, C., Green, M.J., Clark, B.S., Link, B.A., Clarke, J.D., 2013. Mirror-symmetric microtubule assembly and cell interactions drive lumen formation in the zebrafish neural rod. EMBO J. 32, 30–44.

Buckley, C.E., St Johnston, D., 2022. Apical–basal polarity and the control of epithelial form and function. Nat. Rev. Mol. Cell Biol. 1–19.

Cao, L.G., Wang, Y., 1990. Mechanism of the formation of contractile ring in dividing cultured animal cells. II. Cortical movement of microinjected actin filaments. J. Cell Biol. 111, 1905–1911.

Carvalho, C.A., Moreira, S., Ventura, G., Sunkel, C.E., Morais-de-Sá, E., 2015. Aurora A triggers Lgl cortical release during symmetric division to control planar spindle orientation. Curr. Biol. 25, 53–60.

Chen, W., Foss, M., Tseng, K.-F., Zhang, D., 2008. Redundant mechanisms recruit actin into the contractile ring in silkworm spermatocytes. PLoS Biol. 6, e209.

Cheng, N.N., Kirby, C.M., Kemphues, K.J., 1995. Control of cleavage spindle orientation in Caenorhabditis elegans: the role of the genes par-2 and par-3. Genetics 139, 549–559.

Cowan, C.R., Hyman, A.A., 2004. Asymmetric cell division in C. elegans: cortical polarity and spindle positioning. Annu. Rev. Cell Dev. Biol. 20, 427.

Dickinson, D.J., Schwager, F., Pintard, L., Gotta, M., Goldstein, B., 2017. A single-cell biochemistry approach reveals PAR complex dynamics during cell polarization. Dev. Cell 42, 416-434. e11.

Edgar, L.G., Goldstein, B., 2012. Culture and manipulation of embryonic cells, in: Methods in Cell Biology. Elsevier, pp. 151–175.

Etemad-Moghadam, B., Guo, S., Kemphues, K.J., 1995. Asymmetrically distributed PAR-3 protein contributes to cell polarity and spindle alignment in early C. elegans embryos. Cell 83, 743–752.

Etienne-Manneville, S., 2004. Cdc42-the centre of polarity. J. Cell Sci. 117, 1291–1300.

Goehring, N.W., Hoege, C., Grill, S.W., Hyman, A.A., 2011a. PAR proteins diffuse freely across the anterior–posterior boundary in polarized C. elegans embryos. J. Cell Biol. 193, 583–594.

Goehring, N.W., Trong, P.K., Bois, J.S., Chowdhury, D., Nicola, E.M., Hyman, A.A., Grill, S.W., 2011b. Polarization of PAR proteins by advective triggering of a pattern-forming system. Science 334, 1137–1141.

Gotta, M., Abraham, M.C., Ahringer, J., 2001. CDC-42 controls early cell polarity and spindle orientation in C. elegans. Curr. Biol. 11, 482–488.

Gross, P., Kumar, K.V., Goehring, N.W., Bois, J.S., Hoege, C., Jülicher, F., Grill, S.W., 2019. Guiding self-organized pattern formation in cell polarity establishment. Nat. Phys. 15, 293–300.

Guo, S., Kemphues, K.J., 1995. par-1, a gene required for establishing polarity in C. elegans embryos, encodes a putative Ser/Thr kinase that is asymmetrically distributed. Cell 81, 611–620.

Hao, Y., Boyd, L., Seydoux, G., 2006. Stabilization of cell polarity by the C. elegans RING protein PAR-2. Dev. Cell 10, 199–208.

Harris, K.P., Tepass, U., 2010. Cdc42 and vesicle trafficking in polarized cells. Traffic 11, 1272–1279.

Harris, T.J., Peifer, M., 2004. Adherens junction-dependent and-independent steps in the establishment of epithelial cell polarity in Drosophila. J. Cell Biol. 167, 135–147.

Hoege, C., Constantinescu, A.-T., Schwager, A., Goehring, N.W., Kumar, P., Hyman, A.A., 2010. LGL can partition the cortex of one-cell Caenorhabditis elegans embryos into two domains. Curr. Biol. 20, 1296–1303.

Hubatsch, L., Peglion, F., Reich, J.D., Rodrigues, N.T., Hirani, N., Illukkumbura, R., Goehring, N.W., 2019. A cell-size threshold limits cell polarity and asymmetric division potential. Nat. Phys. 15, 1078–1085.

Illukkumbura, R., Hirani, N., Borrego-Pinto, J., Bland, T., Ng, K., Hubatsch, L., McQuade, J., Endres, R., Goehring, N., 2022. Design principles for selective polarization of PAR proteins by cortical flows. bioRxiv.

Kamath, R.S., Ahringer, J., 2003. Genome-wide RNAi screening in Caenorhabditis elegans. Methods 30, 313–321.

Khaliullin, R.N., Green, R.A., Shi, L.Z., Gomez-Cavazos, J.S., Berns, M.W., Desai, A., Oegema, K., 2018. A positive-feedback-based mechanism for constriction rate acceleration during cytokinesis in Caenorhabditis elegans. Elife 7, e36073.

Klinkert, K., Levernier, N., Gross, P., Gentili, C., von Tobel, L., Pierron, M., Busso, C., Herrman, S., Grill, S.W., Kruse, K., 2019. Aurora A depletion reveals centrosome-independent polarization mechanism in Caenorhabditis elegans. Elife 8, e44552.

Klompstra, D., Anderson, D.C., Yeh, J.Y., Zilberman, Y., Nance, J., 2015. An instructive role for C. elegans E-cadherin in translating cell contact cues into cortical polarity. Nat. Cell Biol. 17, 726–735.

Kumfer, K.T., Cook, S.J., Squirrell, J.M., Eliceiri, K.W., Peel, N., O’Connell, K.F., White, J.G., 2010. CGEF-1 and CHIN-1 regulate CDC-42 activity during asymmetric division in the Caenorhabditis elegans embryo. Mol. Biol. Cell 21, 266–277.

Li, D., Mangan, A., Cicchini, L., Margolis, B., Prekeris, R., 2014. FIP 5 phosphorylation during mitosis regulates apical trafficking and lumenogenesis. EMBO Rep. 15, 428–437.

Liang, X., Weberling, A., Hii, C.Y., Zernicka-Goetz, M., Buckley, C.E., 2022. E-cadherin mediates apical membrane initiation site localisation during de novo polarisation of epithelial cavities. EMBO J. e111021.

Lim, Y.W., Wen, F.-L., Shankar, P., Shibata, T., Motegi, F., 2021. A balance between antagonizing PAR proteins specifies the pattern of asymmetric and symmetric divisions in C. elegans embryogenesis. Cell Rep. 36, 109326.

Liu, J., Maduzia, L.L., Shirayama, M., Mello, C.C., 2010. NMY-2 maintains cellular asymmetry and cell boundaries, and promotes a SRC-dependent asymmetric cell division. Dev. Biol. 339, 366–373.

Longhini, K.M., Glotzer, M., 2022. Aurora A and cortical flows promote polarization and cytokinesis by inducing asymmetric ECT-2 accumulation. eLife 11, e83992.

Luján, P., Varsano, G., Rubio, T., Hennrich, M.L., Sachsenheimer, T., Gálvez-Santisteban, M., Martín-Belmonte, F., Gavin, A.-C., Brügger, B., Köhn, M., 2016. PRL-3 disrupts epithelial architecture by altering the post-mitotic midbody position. J. Cell Sci. 129, 4130–4142.

Meitinger, F., Khmelinskii, A., Morlot, S., Kurtulmus, B., Palani, S., Andres-Pons, A., Hub, B., Knop, M., Charvin, G., Pereira, G., 2014. A memory system of negative polarity cues prevents replicative aging. Cell 159, 1056–1069.

Miller, K.E., Kang, P.J., Park, H.-O., 2020. Regulation of Cdc42 for polarized growth in budding yeast. Microb. Cell 7, 175.

Motegi, F., Zonies, S., Hao, Y., Cuenca, A.A., Griffin, E., Seydoux, G., 2011. Microtubules induce self-organization of polarized PAR domains in Caenorhabditis elegans zygotes. Nat. Cell Biol. 13, 1361–1367.

Munro, E., Nance, J., Priess, J.R., 2004. Cortical flows powered by asymmetrical contraction transport PAR proteins to establish and maintain anterior-posterior polarity in the early C. elegans embryo. Dev. Cell 7, 413–424.

Ng, K., Bland, T., Hirani, N., Goehring, N.W., 2022. An analog sensitive allele permits rapid and reversible chemical inhibition of PKC-3 activity in C. elegans. Micropublication Biol. 2022.

Peglion, F., Goehring, N.W., 2019. Switching states: dynamic remodelling of polarity complexes as a toolkit for cell polarization. Curr. Opin. Cell Biol. 60, 121–130.

Pimpale, L.G., Middelkoop, T.C., Mietke, A., Grill, S.W., 2020. Cell lineage-dependent chiral actomyosin flows drive cellular rearrangements in early Caenorhabditis elegans development. Elife 9, e54930.

Pittman, K.J., Skop, A.R., 2012. Anterior PAR proteins function during cytokinesis and maintain DYN-1 at the cleavage furrow in Caenorhabditis elegans. Cytoskeleton 69, 826–839.

Pollarolo, G., Schulz, J.G., Munck, S., Dotti, C.G., 2011. Cytokinesis remnants define first neuronal asymmetry in vivo. Nat. Neurosci. 14, 1525–1533.

Rathbun, L.I., Colicino, E.G., Manikas, J., O’Connell, J., Krishnan, N., Reilly, N.S., Coyne, S., Erdemci-Tandogan, G., Garrastegui, A., Freshour, J., 2020. Cytokinetic bridge triggers de novo lumen formation in vivo. Nat. Commun. 11, 1–12.

Reich, J.D., Hubatsch, L., Illukkumbura, R., Peglion, F., Bland, T., Hirani, N., Goehring, N.W., 2019. Regulated activation of the PAR polarity network ensures a timely and specific response to spatial cues. Curr. Biol. 29, 1911-1923. e5.

Rodrigues, N.T.L., Bland, T., Borrego-Pinto, J., Ng, K., Hirani, N., Gu, Y., Foo, S., Goehring, N.W., 2022. SAIBR: A simple, platform-independent method for spectral autofluorescence correction. Development dev. 200545. https://doi.org/10.1242/dev.200545

Rodriguez, J., Peglion, F., Martin, J., Hubatsch, L., Reich, J., Hirani, N., Gubieda, A.G., Roffey, J., Fernandes, A.R., St Johnston, D., 2017. aPKC cycles between functionally distinct PAR protein assemblies to drive cell polarity. Dev. Cell 42, 400-415. e9.

Rose, L., Gönczy, P., 2014. Polarity establishment, asymmetric division and segregation of fate determinants in early C. elegans embryos. WormBook.

Sailer, A., Anneken, A., Li, Y., Lee, S., Munro, E., 2015. Dynamic opposition of clustered proteins stabilizes cortical polarity in the C. elegans zygote. Dev. Cell 35, 131–142.

Schierenberg, E., 1987. Reversal of cellular polarity and early cell-cell interaction in the embryo of Caenorhabditis elegans. Dev. Biol. 122, 452–463.

Schlüter, M.A., Pfarr, C.S., Pieczynski, J., Whiteman, E.L., Hurd, T.W., Fan, S., Liu, C.-J., Margolis, B., 2009. Trafficking of Crumbs3 during cytokinesis is crucial for lumen formation. Mol. Biol. Cell 20, 4652–4663.

Schonegg, S., Hyman, A.A., 2006. CDC-42 and RHO-1 coordinate acto-myosin contractility and PAR protein localization during polarity establishment in C. elegans embryos. Stiernagle, T., 1999. Maintenance of C. elegans.

Sulston, J.E., Schierenberg, E., White, J.G., Thomson, J.N., 1983. The embryonic cell lineage of the nematode Caenorhabditis elegans. Dev. Biol. 100, 64–119.

Tabuse, Y., Izumi, Y., Piano, F., Kemphues, K.J., Miwa, J., Ohno, S., 1998. Atypical protein kinase C cooperates with PAR-3 to establish embryonic polarity in Caenorhabditis elegans. Development 125, 3607–3614.

Tawk, M., Araya, C., Lyons, D.A., Reugels, A.M., Girdler, G.C., Bayley, P.R., Hyde, D.R., Tada, M., Clarke, J.D., 2007. A mirror-symmetric cell division that orchestrates neuroepithelial morphogenesis. Nature 446, 797–800.

Thielicke, W., Stamhuis, E., 2014. PIVlab–towards user-friendly, affordable and accurate digital particle image velocimetry in MATLAB. J. Open Res. Softw. 2.

Wang, T., Yanger, K., Stanger, B.Z., Cassio, D., Bi, E., 2014. Cytokinesis defines a spatial landmark for hepatocyte polarization and apical lumen formation. J. Cell Sci. 127, 2483–2492.

Watts, J.L., Etemad-Moghadam, B., Guo, S., Boyd, L., Draper, B.W., Mello, C.C., Priess, J.R., Kemphues, K.J., 1996. par-6, a gene involved in the establishment of asymmetry in early C. elegans embryos, mediates the asymmetric localization of PAR-3. Development 122, 3133–3140.

Wirtz-Peitz, F., Nishimura, T., Knoblich, J.A., 2008. Linking cell cycle to asymmetric division: Aurora-A phosphorylates the Par complex to regulate Numb localization. cell 135, 161–173.

Yumura, S., 2001. Myosin II dynamics and cortical flow during contractile ring formation in Dictyostelium cells. J. Cell Biol. 154, 137–146.

Zonies, S., Motegi, F., Hao, Y., Seydoux, G., 2010. Symmetry breaking and polarization of the C. elegans zygote by the polarity protein PAR-2. Development 137, 1669–1677.

