## Supplemental Materials for "Cleavage furrow-directed cortical flows bias mechanochemical pathways for PAR polarization in the *C. elegans* germ lineage"

Table S1

Figures S1-S8

Movie Legends for Movies S1-S4

### Table S1

| Reagent | Description | Source |
|---|---|---|
| Bacteria | | |
| OP50 | E. coli B, ura- | CGC |
| HT115(DE3) | F-, mcrA, mcrB, IN(rrnD-rrnE)1, rnc14::Tn10(DE3 lysogen: lavUV5 promoter-T7 polymerase). | CGC |
| Worms | | |
| BOX241 | *par-6(mib25[par-6::mCherry-LoxP]) I* | Mike Boxem |
| KK1254 | *par-2 (it315[mCherry::par-2]) III* | Ken Kempheus |
| KK1273 | *par-2(it328[GFP::par-2]) III* | CGC/ Ken Kempheus |
| LP162 | *nmy-2(cp13[nmy-2::gfp + LoxP]) I* | CGC |
| LP216 | *par-6(cp45[par-6::mNeonGreen::3xFlag + LoxP unc-119(+) LoxP]) I; unc-119(ed3) III* | Dan Dickinson |
| LP637 | *par-2(cp329[mNG-C1^PAR-2]) III* | Dan Dickinson |
| N2 | Wild type | CGC |
| NWG0076 | *par-6(mib25[par-6::mCherry-LoxP]) I;*<br>*par-2 (it328[gfp::par-2]) III* | This paper |
| NWG0091 | *pkc-3(it309 [gfp::pkc-3]) II;*<br>*par-2 (it315[mCherry::par-2]) III* | This paper |
| NWG0150 | *nmy-2(cp52[nmy-2::mkate2 + LoxP unc-119(+) LoxP]) I;*<br>*par-3(it298 [par-3::gfp]) III* | This paper |
| NWG0189 | *par-3(cp54[mNG::3xFlag::par-3]) III* | Dan Dickinson |
| NWG0192 | *par-2(crk30[par-2(R183-5A)::gfp]*it328) III* | This paper |
| NWG0240 | *par-2(crk41[mNG::par-2(C56S)]*cp329) / sC1(s2023) [dpy-1(s2170) umnIs21] III* | This paper |
| NWG0258 | *par-6(mib25[par-6::mCherry-LoxP]) I;*<br>*par-2 (it328[gfp::par-2]) par-3(crk46[par-3(S950A)]) III* | This paper |
| NWG0259 | *par-3(crk47[mNG::3xFlag::par-3(S950A)]*cp54) III* | This paper |
| NWG0268 | *par-6(cp45[par-6::mNeonGreen::3xFlag + LoxP unc-119(+) LoxP]) I;*<br>*par-2 (it315[mCherry::par-2]) unc-119(ed3) III?* | This paper |
| NWG0283 | *nmy-2(ne3409) par-6(mib25[par-6::mCherry-LoxP]) I;*<br>*par-2(it328[gfp::par-2]) III* | This paper |
| NWG0290 | *par-6(mib25[par-6::mCherry-LoxP]) I;*<br>*lgl-1(crk67[LGL-1::GFP]) X* | (Rodrigues et al., 2022) |

| | | |
|---|---|---|
| NWG0316 | *pkc-3(crk77[I331A,T394A]) II;*<br>*par-2(it328[gfp::par-2]) III* | (Ng et al., 2022) |
| NWG0319 | *nmy-2(ne3409) par-6(mib25[par-6::mCherry-LoxP]) I;*<br>*par-2 (it328[gfp::par-2]) par-3(crk46[par-3(S950A)]) III* | This paper |
| NWG0332 | *par-2 (it315[mCherry::par-2]) III;*<br>*par-1(ax4206) V* | This paper |
| NWG0343 | *par-6(mib25[par-6::mCherry-LoxP]) I;*<br>*par-1(ax4206) V* | This paper |
| NWG0344 | *par-2(crk3[par-2(S241A)::mCherry]*it315)/ sC1(s2023)*<br>*[dpy-1(s2170) umnIs21] III;*<br>*par-1(ax4206) V* | This paper |
| NWG0360 | *par-6(mib25[par-6::mCherry-LoxP]) I;*<br>*par-2(crk96[par-2(S241A)]) III;*<br>*par-1(ax4206) V* | This paper |
| NWG0453 | *par-2 (it315[mCherry::par-2]) par-3(cp54[mNG::3xFlag::par-3]) III* | This paper |
| NWG0458 | *pkc-3(crk77 [I331A,T394A]) II*<br>*par-2 (it315[mCherry::par-2]) par-3(cp54[mNG::3xFlag::par-3]) III* | This paper |
| NWG0473 | *par-2(crk159[mNG::par-2(S241A,GCN4(LI-4mer))*crk104]) /*<br>*sC1(s2023) [dpy-1(s2170) umnIs21] III* | (Illukumbura et al., 2022) |
| OD58 | *unc-119(ed3) III; ltIs38[pAA1; pie-1::GFP::PH(PLC1delta1) +*<br>*unc-119(+)].* | CGC |
| Recombinant DNA - RNAi Clones | | |
| Ahringer Feeding RNAi: *perm-1* | WB Clone: sjj_T01H3.4 | Source BioScience |
| Ahringer Feeding RNAi: *ptr-2* | WB Clone: sjj_C32E8.8 | Source BioScience |
| Recombinant Nucleotides - CRISPR | | |
| PAR-2(C56S) sgRNA #1 | GCTGATCACACAGTGGACAG | |
| PAR-2(C56S) sgRNA #2 | TCGAAAAGCTGATCACACAG | |
| PAR-2(C56S) FWD ODN | GATTGCCAACTCATCGCCAC | |
| PAR-2(C56S) REV ODN | TCCGGCAAAATTGGGGTTTT | |
| PAR-2(C56S) Repair template | TGCATCAACGACGTTCAACAGCCGGTTCGACGCGATTTGAGCT<br>CGGAACTCTTAAGCCCCCTGTGTGAT**CAATTG**TTCGACAGGGTT | |

| | |
|---|---|
| (**MfeI site**) | AGAACATGGAAA |
| PAR-2(R183-5A) sgRNA #1 | CAGTGGGCGACGGCGGTGTG |
| PAR-2(R183-5A) sgRNA #2 | TTGGCGACAGTGGGCGACGG |
| PAR-2(R183-5A) FWD ODN | TCACCGAGCACATTTGACCA |
| PAR-2(R183-5A) REV ODN | AGCTATTCGGGGCGGAAAAA |
| PAR-2(R183-5A) Repair template (**NotI site**) | TCCCATCAAAATCTTTATTTTTTCAGCCCAATCACCCCACAC**GCGGCCGC**TCCACTGTCGCCAAGTGCTCGTCCCGCCAAAAGTTCTCTGAAAATCCCGC |
| PAR-3(S950A) sgRNA #1 | AAACCGACGCTCACAAGCTA |
| PAR-3(S950A) sgRNA #2 | GGAGAGCATTAACCGACCTG |
| PAR-3(S950A) FWD ODN | TGTTGATGACCACGACCCTG |
| PAR-3(S950A) REV ODN | AGGTGGAGCACGTTCAGATG |
| PAR-3(S950A) Repair template (**MspI site**) | TTAAC**CCGG**TTTCAGATATGCTGAATCGCCGTTCTCAGGCAATGGAGAGCATTAACCGACCTGTAGAGAGCATTCTCCGTGGAACGGGGCAAATTCCAACA |

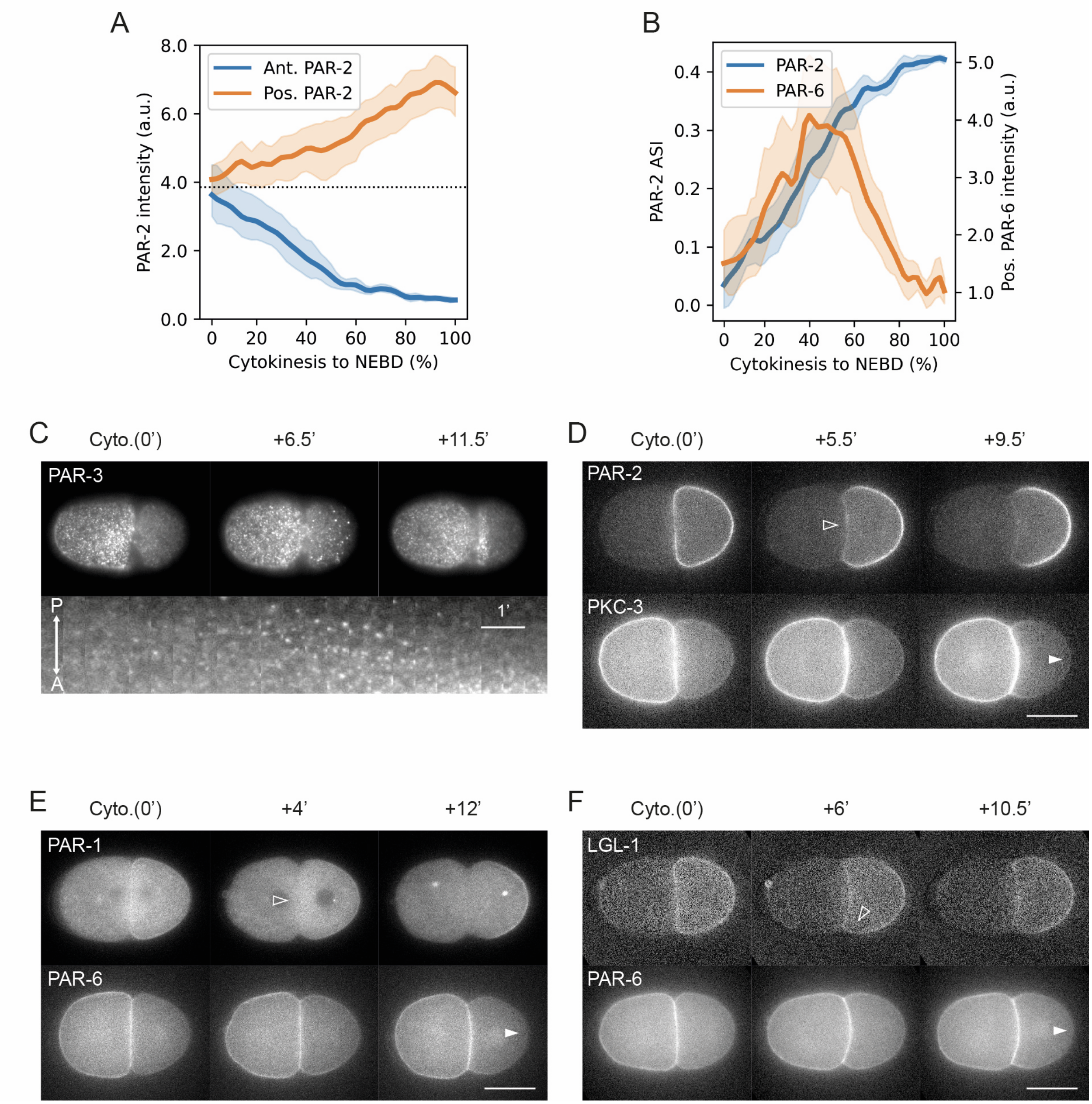
A
Ant. PAR-2
Pos. PAR-2
PAR-2 intensity (a.u.)
Cytokinesis to NEBD (%)
B
PAR-2
PAR-6
PAR-2 ASI
Pos. PAR-6 intensity (a.u.)
Cytokinesis to NEBD (%)
C
Cyto.(0')
+6.5'
+11.5'
PAR-3
P
A
1'
D
Cyto.(0')
+5.5'
+9.5'
PAR-2
PKC-3
E
Cyto.(0')
+4'
+12'
PAR-1
PAR-6
F
Cyto.(0')
+6'
+10.5'
LGL-1
PAR-6

**Figure S1. Polarization of pPARs precedes aPAR segregation in P1.**

(A, B) Further quantification of embryos expressing both mCherry::PAR-2 and PAR-6::mNG (NWG0268) during polarity establishment in P1, corresponding to Figure 1. (A) PAR-2 levels increase in the posterior as PAR-2 in the anterior is cleared, suggesting coalescence towards the posterior. (B) PAR-2 develops an asymmetry early towards the posterior despite an initial corresponding rise in PAR-6 levels in the posterior. ASI = asymmetry index.

(C) Cortical imaging of mNG::PAR-3 (NWG0189) (n=8) through the cell cycle of P1. Bottom, a kymograph of PAR-3 localization in P1 for the same embryo.

(D-F) Time series of midsection confocal images of an embryo expressing both mCherry::PAR-2 and GFP::PKC-3 (NWG0091) (n=3) (D), PAR-1::GFP and PAR-6::mCherry (NWG0343) (n=3) (E), and LGL-1::GFP and PAR-6::mCherry (NWG0290) (n=3) (F), during polarity establishment in P1. Open arrowheads indicate when notable pPAR clearance can be seen, and closed arrowheads indicate when segregation of aPARs can be seen. Scale bar, 20μm.

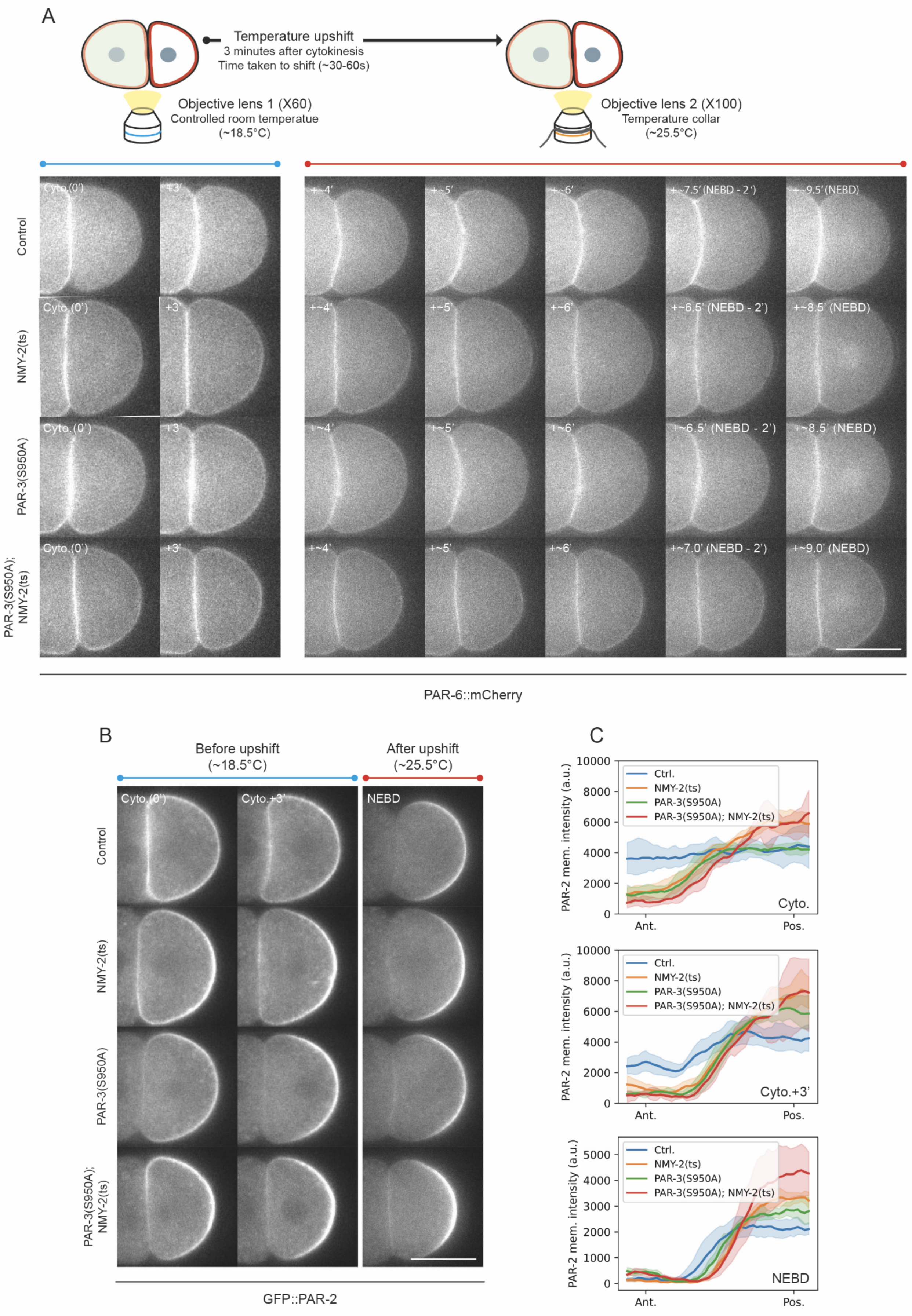
A
Temperature upshift
3 minutes after cytokinesis
Time taken to shift (~30-60s)
Objective lens 1 (X60)
Controlled room temperatue
(~18.5°C)
Objective lens 2 (X100)
Temperature collar
(~25.5°C)
Control
NMY-2(ts)
PAR-3(S950A)
PAR-3(S950A); NMY-2(ts)
Cyto.(0')
+3'
+~4'
+~5'
+~6'
+~7.5' (NEBD - 2')
+~9.5' (NEBD)
+~6.5' (NEBD - 2')
+~8.5' (NEBD)
+~7.0' (NEBD - 2')
+~9.0' (NEBD)
PAR-6::mCherry
B
Before upshift
(~18.5°C)
After upshift
(~25.5°C)
Cyto.(0')
Cyto.+3'
NEBD
GFP::PAR-2
C
Ctrl.
NMY-2(ts)
PAR-3(S950A)
PAR-3(S950A); NMY-2(ts)
PAR-2 mem. intensity (a.u.)
Cyto.
Cyto.+3'
NEBD
Ant.
Pos.
0
2000
4000
6000
8000
10000
1000
3000
5000

**Figure S2. PAR polarization in P1 cells in a *nmy-2(ts)* and *par-3(S950A)* background.**
(A) Top, a schematic illustrating the setup used to achieve acute temperature upshift. Bottom, a time series of midsection confocal images of embryos expressing GFP::PAR-2 (not shown) and PAR-6::mCherry in a wild-type (Ctrl.; NWG0076), *nmy-2(ts)* (NWG0283), *par-3(S950A)* (NWG0258), or a double *nmy-2(ts); par-3(S950A)* background (NWG0319) at indicated times. Scale bar, 20μm. Sample sizes are identical to Figure 2. Note that the bulging of the AB-P1 cell contact, which depends on NMY-2 activity, is lost in *nmy-2(ts)* embryos when upshifted to the restrictive temperature, consistent with efficient disruption of NMY-2 activity.
(B) Time series of midsection confocal images of the same embryos in (A) but showing GFP::PAR-2 instead. Sample sizes are identical to Figure 2. Scale bar, 20μm.
(C) Quantification of PAR-2 membrane profiles corresponding to conditions shown in (B). Note that due to changes in objectives for the temperature shift, intensity values between Cyto and Cyto+3’ are not directly comparable to those at NEBD. Mean and 95% confidence interval (bootstrapped) indicated.

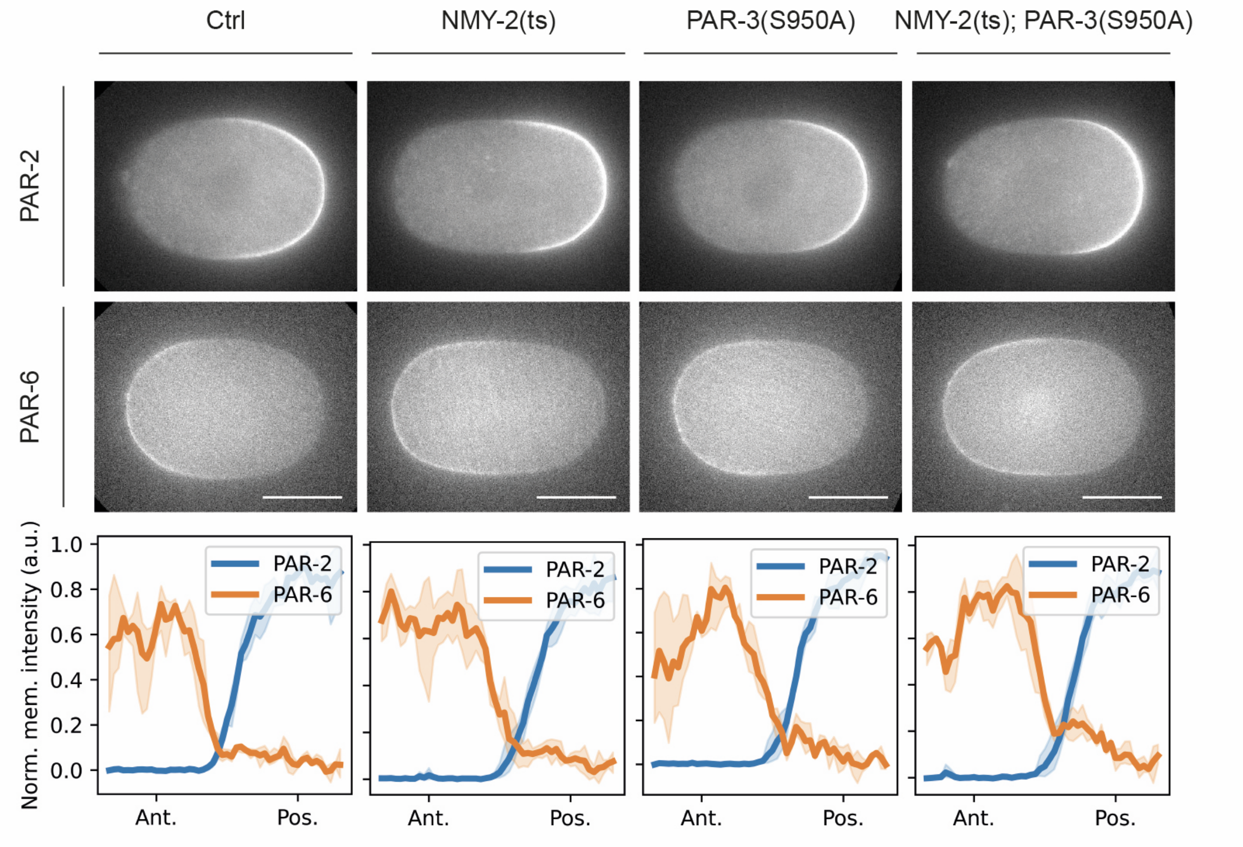
Ctrl
NMY-2(ts)
PAR-3(S950A)
NMY-2(ts); PAR-3(S950A)
PAR-2
PAR-6
Norm. mem. intensity (a.u.)
1.0
0.8
0.6
0.4
0.2
0.0
PAR-2
PAR-6
Ant.
Pos.

**Figure S3. PAR polarization in the zygote is normal in a combination of *nmy-2(ts)* and *par-3(S950A)* backgrounds under permissive temperature (18.5ºC).**

Top, representative midsection confocal images of embryos expressing GFP::PAR-2 and PAR-6::mCherry in a wild-type (Ctrl.; NWG0076) (n=3), *nmy-2(ts)* (NWG0283) (n=3), *par-3(S950A)* (NWG0258) (n=4), or a double *nmy-2(ts); par-3(S950A)* background (NWG0319) (n=3) during NEBD.

Bottom, quantification of normalized membrane profiles of embryos corresponding to genotypes indicated above. Mean and 95% confidence interval (bootstrapped) indicated. Scale bar, 20μm.

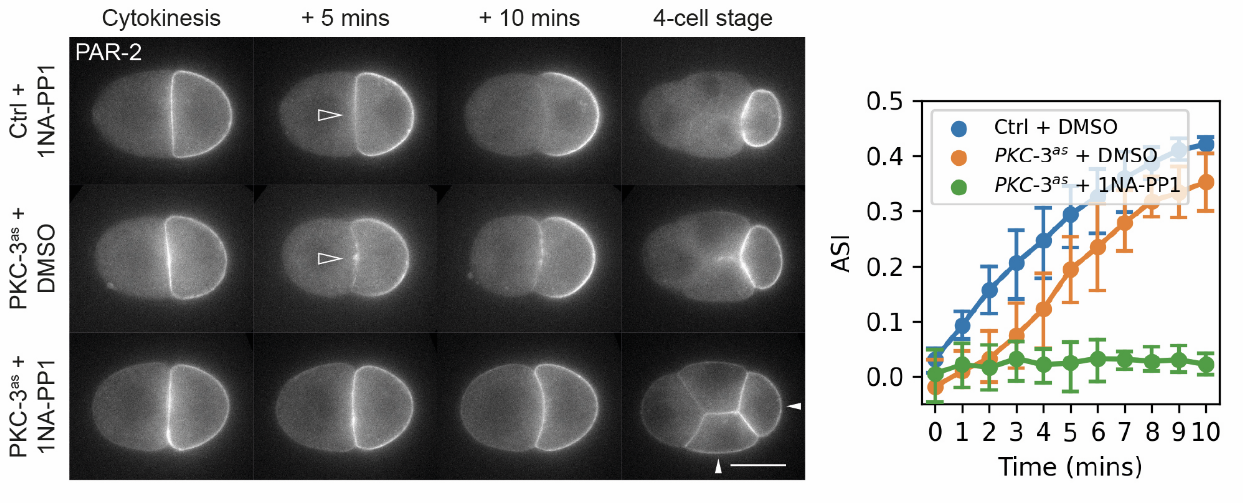
Cytokinesis
+ 5 mins
+ 10 mins
4-cell stage
PAR-2
Ctrl + 1NA-PP1
PKC-3as + DMSO
PKC-3as + 1NA-PP1
Ctrl + DMSO
PKC-3as + DMSO
PKC-3as + 1NA-PP1
ASI
Time (mins)
0.0
0.1
0.2
0.3
0.4
0.5
0 1 2 3 4 5 6 7 8 9 10

**Figure S4. PKC-3 activity is required for PAR-2 polarization in P1**

Left, a time series of midsection confocal images of embryos expressing GFP::PAR-2 in either a PKC-3(WT) (KK1273) (n=5), or PKC-$3^{AS}$ background (NWG0316), treated with either DMSO (n=5) or 20μM of the ATP analog 1NA-PP1 (Calbiochem) (n=5). Open arrowheads indicate early PAR-2 clearance from the cell anterior, and closed arrowheads indicate symmetric sized daughter cells. Scale bar, 20μm.

Right, quantification of asymmetry index (ASI) for times indicated for the corresponding conditions. Mean and 95% confidence interval (bootstrapped) indicated.

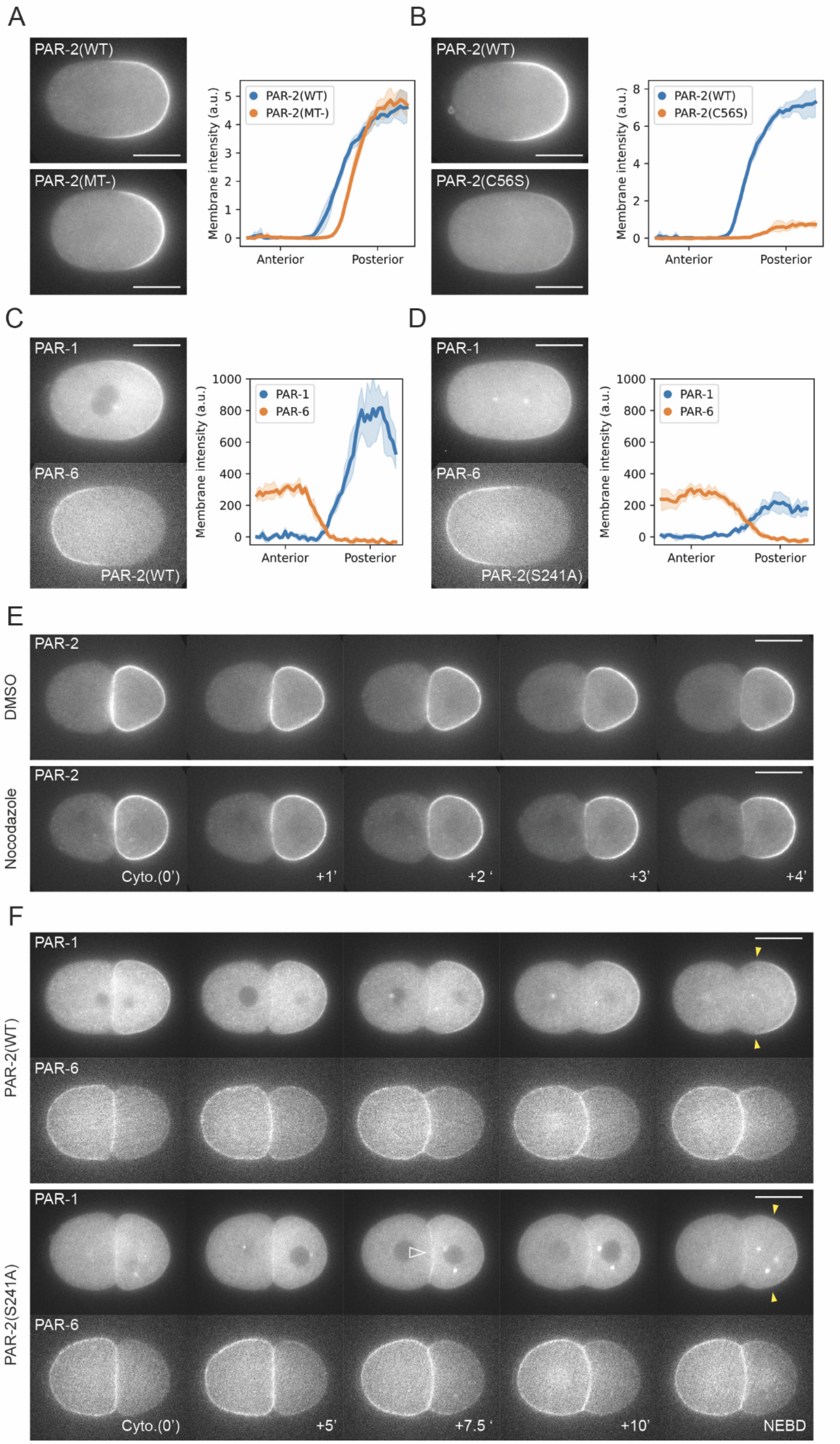
A
PAR-2(WT)
PAR-2(MT-)
Membrane intensity (a.u.)
Anterior
Posterior
B
PAR-2(WT)
PAR-2(C56S)
C
PAR-1
PAR-6
PAR-2(WT)
D
PAR-2(S241A)
E
PAR-2
DMSO
Nocodazole
Cyto.(0')
+1'
+2 '
+3'
+4'
F
+5'
+7.5 '
+10'
NEBD

**Figure S5. PAR polarization in conditions in which PAR-2 self-organization is perturbed in the zygote and P1**

(A-D) Midsection confocal images of zygotes during NEBD and corresponding membrane profile quantifications for indicated conditions. Sample sizes: (A) PAR-2(WT) (n=5), PAR-2(MT-) (n=7), (B) PAR-2(WT) (n=8), PAR-2(C56S) (n=5), (C) PAR-2(WT) (n=4), (D) PAR-2(S241A) (n=7). Mean and 95% confidence interval (bootstrapped) indicated.

(E) A time series of midsection confocal images of embryos expressing GFP::PAR-2 (KK1273), treated with either DMSO (n=2) or nocodazole (10 μg/ml) (n=2).

(F) A time series of midsection confocal images of embryos expressing both PAR-1::GFP and PAR-6::mCherry in either a wild type background (NWG0343) (n=4) or PAR-2(S241A) background (NWG0360) (n=4). Open arrowhead indicates late clearance of PAR-1 from the anterior in *par-2(S241A)* embryos relative to *par-2(WT)* embryos. Closed yellow arrowheads roughly indicate the extent of the PAR-1 domain.

Scale bar, 20μm.

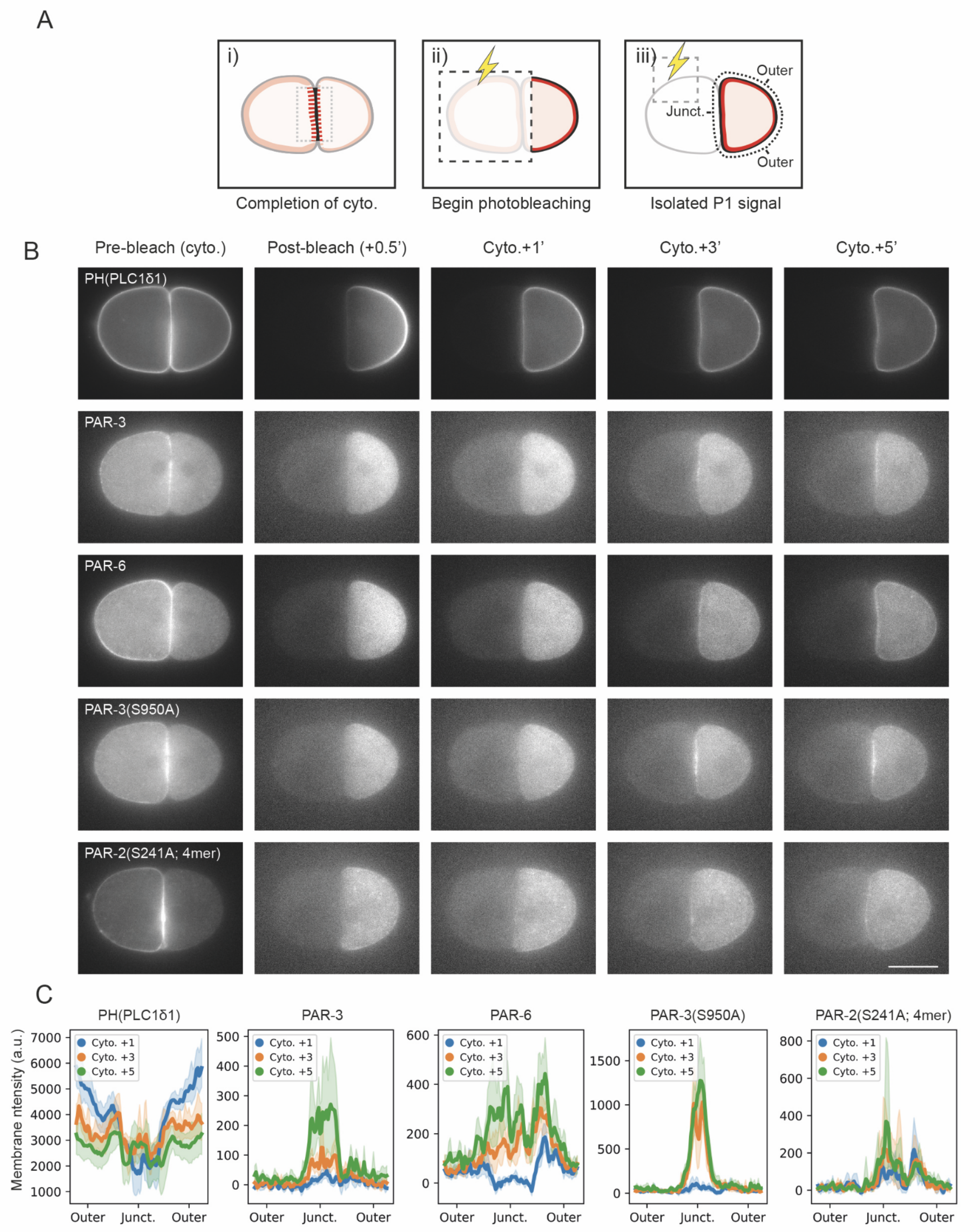
A
i)
Completion of cyto.
ii)
Begin photobleaching
iii)
Outer
Junct.
Outer
Isolated P1 signal
B
Pre-bleach (cyto.)
Post-bleach (+0.5')
Cyto.+1'
Cyto.+3'
Cyto.+5'
PH(PLC1δ1)
PAR-3
PAR-6
PAR-3(S950A)
PAR-2(S241A; 4mer)
C
PH(PLC1δ1)
PAR-3
PAR-6
PAR-3(S950A)
PAR-2(S241A; 4mer)
Membrane ntensity (a.u.)
Cyto. +1
Cyto. +3
Cyto. +5
Outer
Junct.
Outer

**Figure S6. Selective photobleaching of AB reveals an early aPAR bias towards the nascent cell contact of P1.**

(A) Schematic illustrating the workflow used to eliminate confounding AB fluorescence signals at the AB-P1 contact site. (i) Initially, after completion of P0 cytokinesis, aPAR fluorescence signal at the cell-cell contact (dotted box) could come from either AB or P1 cell. (ii) 30 seconds after P0 cytokinesis, a region encompassing the entire AB cell through to the anterior edge of P1, which includes the AB-P1 contact, was bleached (dotted box). (iii) Following, we observed fluorescence recovery in P1 whilst continually bleaching a region of AB, to further ensure that no fluorescence pool remains in AB that could contribute to the contact site. Consequently, signals at the cell contact should come entirely from P1.

(B) Time series of midsection confocal images of embryos expressing GFP::PH(PLC1δ1) (OD58) (n=6), mNG::PAR-3 (NWG0189) (n=7), PAR-6::mNG (LP216) (n=7), mNG::PAR-3(S950A) (NWG0259) (n=6), and mNG::PAR-2 (S241A; 4mer) (NWG0473) (n=7) subjected to the photobleaching assay depicted in (A). Scale bar, 20μm.

(C) Quantification of membrane profiles (as shown in the dotted lines in (A)(iii)) for indicated time points for embryos with conditions corresponding to (B). Mean and 95% confidence interval (bootstrapped) indicated.

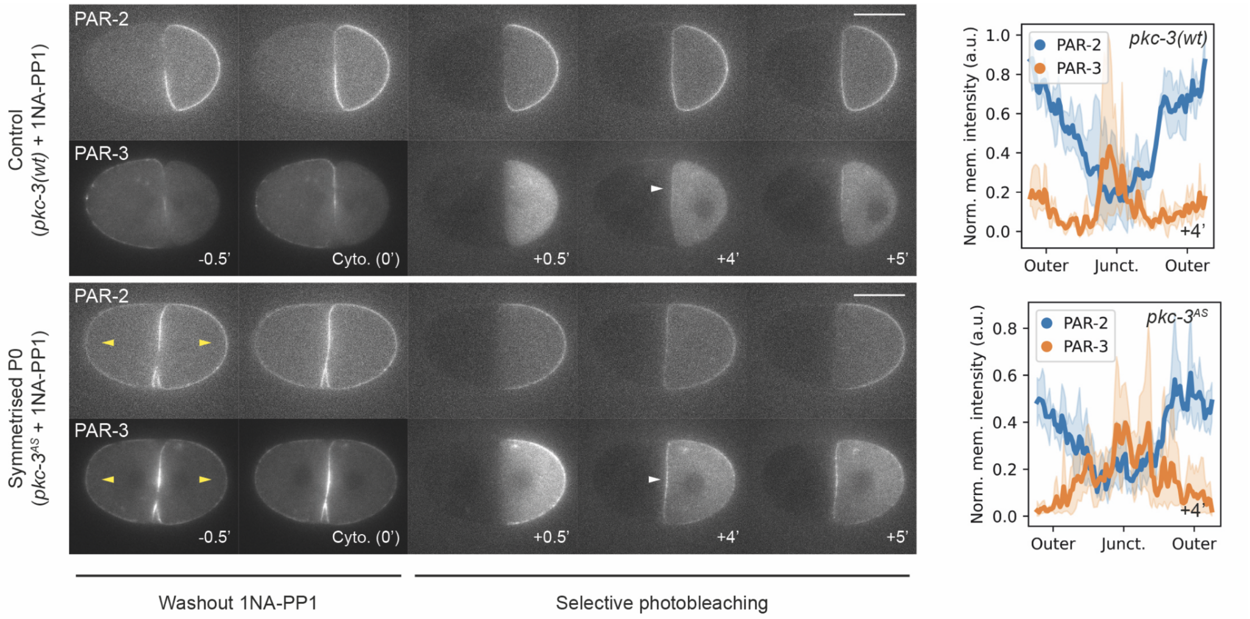

Control
(pkc-3(wt) + 1NA-PP1)
PAR-2
PAR-3
-0.5'
Cyto. (0')
+0.5'
+4'
+5'
Symmetrised P0
(pkc-3AS + 1NA-PP1)
PAR-2
PAR-3
-0.5'
Cyto. (0')
+0.5'
+4'
+5'
Washout 1NA-PP1
Selective photobleaching
pkc-3(wt)
PAR-2
PAR-3
Norm. mem. intensity (a.u.)
1.0
0.8
0.6
0.4
0.2
0.0
+4'
Outer
Junct.
Outer
pkc-3AS
PAR-2
PAR-3
Norm. mem. intensity (a.u.)
0.8
0.6
0.4
0.2
0.0
+4'
Outer
Junct.
Outer

**Figure S7. Activation of PAR activity during P0 furrow closure is sufficient to generate mirror-symmetric PAR polarity.**

Left, a time series of midsection confocal images of embryos at indicated time points expressing mCherry::PAR-2 and mNG::PAR-3 in either PKC-3(WT) (NWG0453) or PKC-$3^{AS}$ (NWG0458) backgrounds, when the zygote is treated with 100μM 1NA-PP1 and washed out during furrow formation, and subjected to the photobleaching protocol depicted in Figure S6A. Yellow arrowheads indicate symmetric inheritance of PAR-3 and PAR-2 in 2-cell embryos following PKC-3 inhibition in the zygote. White arrowheads indicate early anterior enrichment of PAR-3 at the nascent cell contact.

Right, quantification of membrane profiles (as shown in dotted lines in Figure S6Aiii) for the corresponding conditions 4 minutes after completion of cytokinesis. Mean and 95% confidence interval (computed by bootstrapping) indicated.

Scale bars, 20μm.

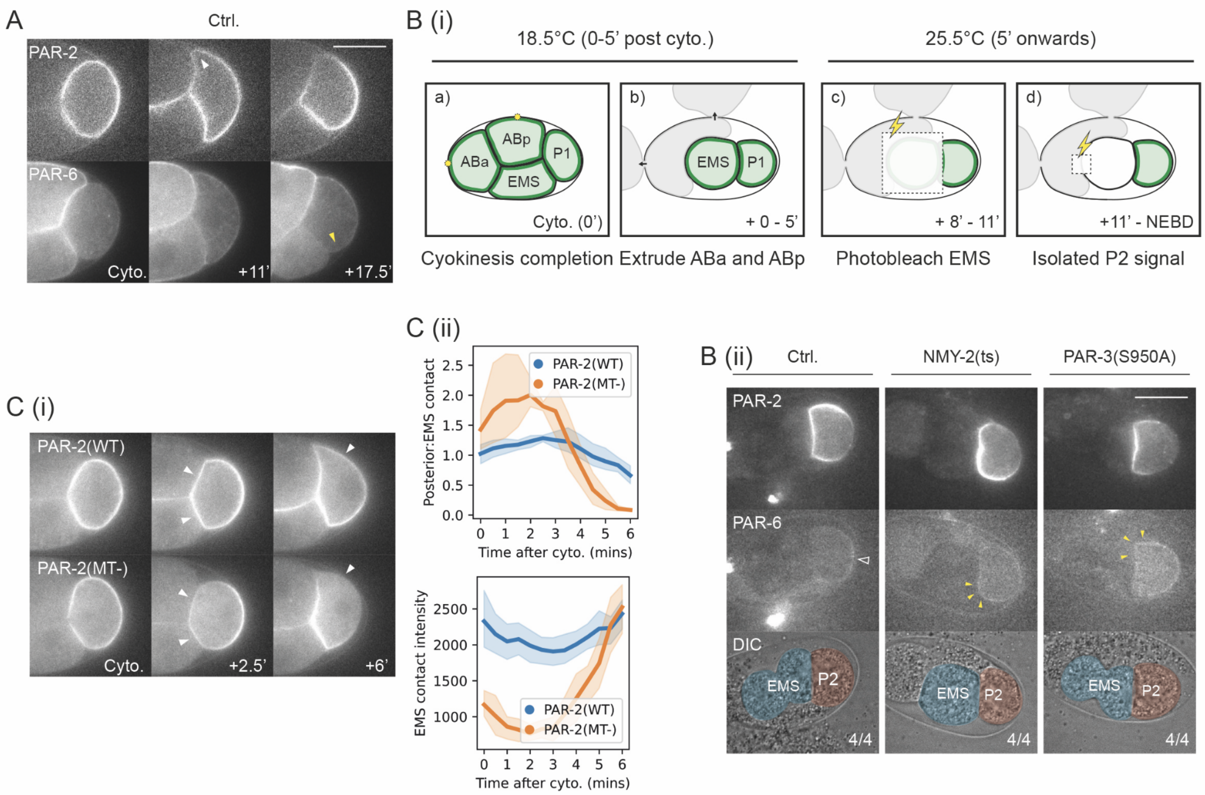
A
Ctrl.
PAR-2
PAR-6
Cyto.
+11'
+17.5'
B (i)
18.5°C (0-5' post cyto.)
25.5°C (5' onwards)
a)
ABa
ABp
P1
EMS
Cyto. (0')
b)
EMS
P1
+ 0 - 5'
c)
+ 8' - 11'
d)
+11' - NEBD
Cyokinesis completion
Extrude ABa and ABp
Photobleach EMS
Isolated P2 signal
C (i)
PAR-2(WT)
PAR-2(MT-)
Cyto.
+2.5'
+6'
C (ii)
Posterior:EMS contact
PAR-2(WT)
PAR-2(MT-)
Time after cyto. (mins)
EMS contact intensity
Time after cyto. (mins)
B (ii)
Ctrl.
NMY-2(ts)
PAR-3(S950A)
PAR-2
PAR-6
DIC
EMS
P2
4/4

**Figure S8. The pattern and mechanisms of polarization are similar in P2 and P1**

(A) Time series of midsection confocal images of embryos expressing both mCherry::PAR-2 and PAR-6::mNG (NWG0268) (n=6) during polarity establishment in P2 at indicated time points. Closed white arrowheads indicate clearance of PAR-2, whereas closed yellow arrowheads indicate clearance of PAR-6. Scale bar, 20μm.

(B)(i) Schematic illustrating the setup used to visualize localization of aPARs in P2. Yellow circles in (a) indicate ablation sites used to lyse ABa and ABp to extrude contents. Dotted boxes in (c) and (d) indicate the regions for photobleaching as in Figure S6A. A large region encompassing EMS and EMS-P1 contact was bleached once (c), and then a smaller region at the anterior of EMS is continuously bleached to ensure no recovery of the fluorescence pool in EMS (d). The protocol used to upshift is identical to Fig S2.

(B)(ii) Midsection confocal images of embryos expressing both GFP::PAR-2 and PAR-6::mCherry in a wildtype (NWG0076), *nmy-2(ts)* (NWG0283), or a *par-3(S950A)* background (NWG0258) subject to the protocol indicated in (B)(i). Open white arrowheads indicate PAR-6 polarization. Closed yellow arrowheads indicate ectopic PAR-6 localization towards the EMS contact, overlapping with the PAR-2 domain. Scale bar, 20μm.

(C)(i) A time series of midsection confocal images of embryos expressing GFP::PAR-2 (KK1273) (n=6) or GFP::PAR-2(R183-5A; MT-) (NWG0192) (n=4). White arrowheads indicate an initial reduction in PAR-2 levels at the anterior of P2 (EMS contact; see 2.5’), before reversing towards a reduction at the posterior pole (see 6’). Scale bar, 20μm.

(C)(ii) Top, quantification of the ratio of PAR-2 membrane signal at the posterior pole to membrane signal at the EMS-P2 contact for conditions corresponding to (C)(i). Bottom, same as top, but for PAR-2 membrane signal.

### Movie S1: Polarization of PAR-2 and PAR-6 in P1 cells

Midsection confocal movie of an embryo expressing both endogenous mCherry::PAR-2 and PAR-6::mNG from cytokinesis to NEBD of P1.

### Movie S2: HiLo imaging of cortical PAR-3 during P0 cytokinesis

HiLo movie of embryos expressing mNG::PAR-3 or mNG::PAR-3(S950A) during P0 cytokinesis. Movies were processed with bleach-correction (histogram matching) and background subtraction. Movement of PAR-3 clusters into the cleavage furrow was highlighted using a rolling temporal-color code with a 7 frame projection, LUT=CET-L18.

### Movie S3: Photobleaching assay to reveal fluorescence signal at cell contacts

Midsection confocal movie of P1 cells that undergo selective photobleaching of AB. Note that the intensity range is renormalized following photobleaching (at time 0.5').

### Movie S4: Inducing mirror-symmetry PAR polarity in 2-cell stage embryos through late activation of PKC-3 activity during furrow closure

Midsection confocal movie of embryos expressing both mCherry::PAR-2 and mNG::PAR-3 in either wildtype (top) or *pkc-3*$^{AS}$ background (bottom). 100μM 1NA-PP1 was washed out following treatment in the zygote.
